# Pigmentation markers enable efficient co-CRISPR selection in the pantry moth *Plodia interpunctella*

**DOI:** 10.64898/2026.11.13.760159

**Authors:** Christa Heryanto, Leah F. Rosin

## Abstract

CRISPR co-targeting (co-CRISPR) of pigment genes facilitates the identification and recovery of genome-edited individuals in model organisms. Using *Plodia interpunctella*, an emerging lepidopteran model, we characterized mutant phenotypes of eight genes involved in the melanin and ommochrome biosynthesis pathways (*black, ebony, yellow, tan*, v*ermilion, kynurenine 3-monooxygenase/cinnabar (kmo/cn), scarlet, and white*) to evaluate their suitability as co-CRISPR markers. Disruption of *black*, *ebony*, and *yellow* altered larval sclerite, pupal cuticle, and adult wing coloration. All four ommochrome pathway genes studied contributed to larval cuticle and adult eye pigmentation, whereas only *vermilion* and *kmo/cn* disruption produced visible phenotypes in the adult wings. Sequencing revealed low apparent mosaicism in some *yellow* and *tan* G_0_ individuals, along with recurrent deletions at the *yellow* and *tan* target sites consistent with possible microhomology-mediated end joining repair. Using *white* or *scarlet* as co-CRISPR selection markers, we identified G_0_ individuals carrying a secondary mutation in the *yellow* gene with a positive predictive value of 100% by scoring its knockout phenotype. Extending this approach to six meiosis-related genes without visibly scorable phenotypes, selection based on the *white* mutant phenotype identified sequence-confirmed mutations at the corresponding gene-of-interest in 191 of 206 G_0_ individuals (93%). Together, these findings establish *white* and *scarlet* as effective co-CRISPR selection markers and provide a practical strategy for facilitating functional genetic studies in *Plodia*.

**GRAPHICAL ABSTRACT:** 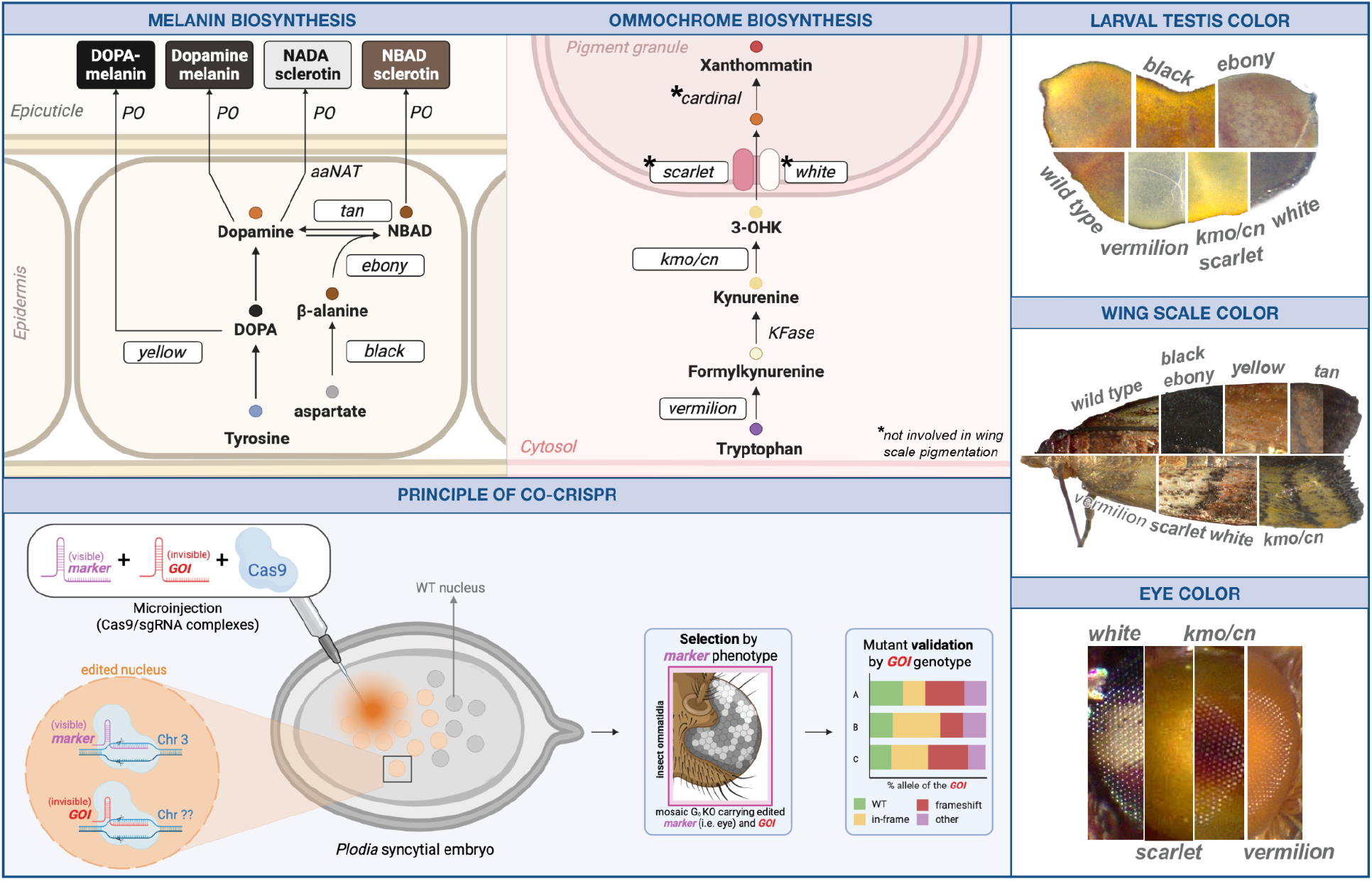

## INTRODUCTION

Understanding how genes control development, physiology, and reproduction requires tools that can reliably disrupt gene function. In moths and butterflies (Lepidoptera), such studies have historically been complicated by the variable effectiveness of RNA interference (RNAi), partly due to RNAi-degrading nucleases (Terenius et al. 2011; Kolliopoulou and Swevers 2014). The introduction of CRISPR/Cas9-mediated genome editing (CRISPR) in Lepidoptera, first demonstrated in the silkworm *Bombyx mori*, has substantially expanded opportunities for functional genetic research in these emerging model systems (Wang et al. 2013; Wei et al. 2014). Although RNAi remains useful for tissue- or stage-specific gene knockdown through approaches such as *in vivo* electroporation of small-interfering RNAs (siRNAs) (Fujiwara and Nishikawa 2016; Okude et al. 2017; Hanly et al. 2023; Livraghi et al. 2025; Loh et al. 2025), embryonic microinjection of CRISPR/Cas9 components remains the gold standard for generating somatic and heritable gene knockouts (KOs) across a wide range of lepidopteran species (Li et al. 2015; Zhang et al. 2017b; You et al. 2019; Shirai et al. 2021; Wang et al. 2021; Xu et al. 2021; Cheng et al. 2023; Pearce et al. 2026).

The pantry moth *Plodia interpunctella* (*Plodia, Pi*) has emerged as a versatile model system for functional genetic research in Lepidoptera. As a silk-producing species, *Plodia* provides opportunities to investigate the genetic regulation of silk gland development and the molecular architecture of silk proteins, with applications in biomaterials research (Alqassar et al. 2025; Eccles et al. 2025; Shodja et al. 2025; Eccles et al. 2026). Its chromosome number, which is more comparable to that of humans than other insect models like *Drosophila*, has also motivated its use in studies of meiosis, chromosome segregation, recombination, and reproduction (Hockens et al. 2024; Lee et al. 2026). Importantly, several genetic tools have already been established in *Plodia*, including CRISPR/Cas9-mediated genome editing (Heryanto et al. 2022a; Shirk et al. 2023; Shodja et al. 2026) and stable transgenesis using *piggyBac* and *Minos* transposable elements (Heryanto et al. 2022b; Shodja et al. 2025). Together with its small size and rapid generation time, these advances make *Plodia* an increasingly accessible system for studying gene function.

Despite these advances, identifying successfully edited individuals remains a major bottleneck in CRISPR-based research. Many genetic modifications produce no immediately visible phenotype, making it difficult to distinguish edited individuals from unedited ones. Researchers must therefore rely on laborious molecular screening and repeated breeding to identify individuals carrying the desired mutations and establish stable mutant lines (Ewen-Campen and Perrimon 2018). The co-CRISPR method, first developed in *C. elegans* (Kim et al. 2014), has reduced this screening burden. This approach simultaneously targets two genes: a gene of interest (GOI) and a marker gene whose disruption produces a readily recognizable phenotype. Since simultaneous editing at both loci occurs at a higher-than-random frequency, and G_0_ individuals displaying the visible marker phenotype are substantially more likely to also carry edits in the GOI, the visible phenotype can help prioritize individuals for subsequent molecular screening and breeding. Co-CRISPR has been effectively demonstrated in the fruitfly *Drosophila melanogaster* and in the fall armyworm *Spodoptera frugiperda*, where the pigment genes *white* and *kynurenine 3-monooxygenase/cinnabar (kmo/cn)*, respectively, served as visible co-CRISPR markers to rapidly identify and enrich GOI mutant selection (Ge et al. 2016). Such approaches offer the potential to reduce the time and resources required to recover desired mutations, particularly in species where genetic screening remains challenging.

Genes involved in insect pigmentation are especially promising candidates for visible CRISPR markers because their disruption can produce easily recognizable changes in coloration. Two major pigmentation pathways, melanin and ommochrome biosynthesis, contribute to the diverse colors and patterns observed in Lepidoptera. In the melanin pathway, *yellow* promotes dark pigment formation, whereas *black* and *ebony* contribute to the production of lighter-colored compounds involved in cuticle hardening. The gene *tan* counteracts *ebony* by converting its product, N-β-alanyldopamine (NBAD), back into dopamine, helping regulate the balance between these pigmentation pathways (Wittkopp et al. 2002; Sugumaran and Barek 2016; Kuwalekar et al. 2020). Beyond their roles in cuticle pigmentation and hardening, melanin pathway genes also contribute to wing scale coloration during pupal development, when pigments are progressively deposited to produce yellow, brown, and black coloration (Koch et al. 1998; Wittkopp and Beldade 2009; Zhang et al. 2017b; Kuwalekar et al. 2020). Disruption of melanin pathway genes, such as *ebony* (Bi et al. 2019; Xu et al. 2021; Inoue et al. 2023; Luo et al. 2025; Su et al. 2025), *tan* (Inoue et al. 2023), and *yellow* (Nie et al. 2021; Wang et al. 2021; Tomihara et al. 2022) produces distinct pigmentation phenotypes in adult body, wings, or scales.

The ommochrome pathway also contributes to insect pigmentation, producing yellow, orange, and red pigments in the eyes and, in some Lepidoptera, wing scales (Reed and Nagy 2005; Ferguson and Jiggins 2009). Ommochrome biosynthesis begins with the conversion of L-tryptophan to formylkynurenine by Vermilion, followed by its conversion to kynurenine by kynurenine formamidase (KFase). KMO/Cinnabar then converts kynurenine to 3-hydroxykynurenine (3-OHK), which is transported into pigment granules by the White-Scarlet transporter complex and subsequently converted into xanthommatin and related pigments (Mackenzie et al. 1999; Quan et al. 2002). Ommochrome-derived pigments contribute to wing coloration in several butterflies, including *Heliconius*, *Araschnia levana*, and *Papilio xuthus* (Ishizaki and Umebachi 1984; Gilbert et al. 1988; Koch 1991). Disruption of genes associated with ommochrome biosynthesis and transport, including *white*, *scarlet*, and *kmo/cn*, has produced visible pigmentation phenotypes in several lepidopteran species (Hong et al. 2020; Xu et al. 2020; Heryanto et al. 2022a; Shirk et al. 2023; Shodja et al. 2026). However, the suitability of these genes as visible co-CRISPR markers in *Plodia* remains largely unexplored. Identifying markers that produce readily distinguishable phenotypes without substantially compromising survival, development, or reproduction is essential for facilitating mutant selection and establishing stable edited lines.

In this study, we investigate eight genes associated with the melanin and ommochrome pigmentation pathways in *P. interpunctella*. Specifically, we identify the melanin pathway genes *black, ebony, tan, and yellow*, and the ommochrome pathway genes *vermilion, kmo/cn*, *white*, and *scarlet* in the pantry moth *Plodia interpunctella*. We then characterize the visible phenotypes and fitness effects resulting from the loss-of-function (LOF) of these genes across *Plodia* life stages and subsequent generations. Finally, we evaluate gene candidates for their suitability as visible markers for CRISPR co-conversion experiments. By identifying pigmentation genes that produce readily detectable phenotypes with limited fitness costs, this work will facilitate more efficient identification and recovery of CRISPR-edited individuals in *Plodia*, supporting the continued development of this species as a model for functional genetics.

## RESULTS

### Melanin and ommochrome pathway genes are conserved in Lepidoptera and expressed in testes

To establish a foundation for pigment-based co-CRISPR selection in *Plodia interpunctella* (*Plodia, Pi*), we first identified and characterized orthologs of conserved melanin and ommochrome pathway genes across Lepidoptera **(Figure 1A)** (Massey and Wittkopp 2016; Zhang et al. 2017a; Kuwalekar et al. 2020). We compared the amino acid (AA) sequences of the pigmentation genes *black*, *ebony*, *tan*, *yellow*, *vermilion*, *kmo/cn*, *white*, and *scarlet* across five well-characterized lepidopteran species: *Plutella xylostella (Px), Bombyx mori (Bm), Spodoptera frugiperda (Sf), Galleria mellonella (Gm),* and *Manduca sexta (Ms*), along with *Drosophila melanogaster (Dm)* as an outgroup **(Figure 1B, 1C; Suppl. Table S1)**. With the exception of *yellow*, all genes exhibited high AA sequence conservation among Lepidoptera (∼80-90% similarity; see methods), but only moderate conservation with the *Dm* outgroup (∼60-80% similarity). The *yellow* gene, by contrast, showed more uniform AA sequence similarity across both groups (∼70-80% similarity). These sequence comparisons enabled the identification of Plodia pigmentation gene orthologs and informed the selection of conserved coding regions for sgRNA design in subsequent CRISPR/Cas9 knockout experiments.

**Figure 1.**
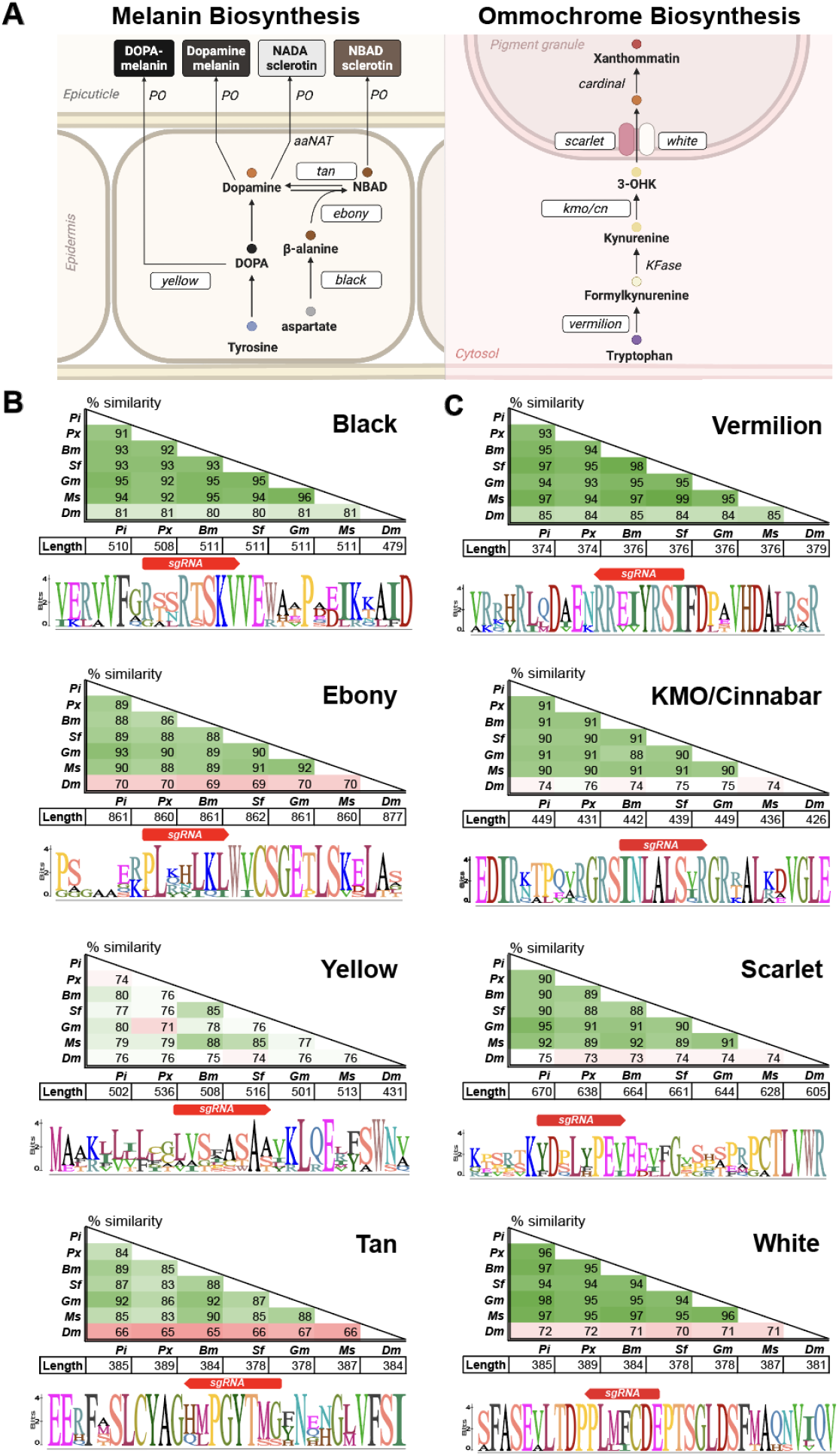
Canonical melanin and ommochrome pathways are well conserved in Lepidoptera. (A) Simplified melanin and ommochrome biosynthesis pathways in insects, adapted from Liao et al. 2018. Genes in the present study are in white boxes. The schematic was created using Biorender. (B) Pairwise comparison of amino acid sequences of Black, Ebony, Yellow, and Tan across representative lepidopteran insects and the outgroup (*Drosophila melanogaster*). (C) Pairwise comparison of amino acid sequences of Vermilion, Kynurenine 3-monooxygenase/Cinnabar (KMO/Cn), Scarlet, and White across representative lepidopteran insects and the outgroup. For (B) and (C), BLOSUM90 amino acid substitution in seven insect species is shown as a similarity matrix (top). Red-white-green shading represents increasing similarity. Under the matrix are the sequence lengths to show that the aligned sequences are of comparable size (middle). The amino acids in the proximity of the KO target coding region used in this study are conserved (bottom). *Pi: Plodia interpunctella, Px: Plutella xylostella, Bm: Bombyx mori, Sf: Spodoptera frugiperda, Gm: Galleria mellonella, Ms: Manduca sexta, Dm: Drosophila melanogaster*.

Given that CRISPR/Cas9 microinjection into early embryos edits both somatic and germline cells, we investigated whether pigmentation phenotypes in the testes could serve as a visible marker of CRISPR activity in gonadal tissues and potentially enrich for individuals carrying germline edits, thereby facilitating co-CRISPR selection. Supporting this idea, prior studies demonstrated that the ommochrome genes *white* and *scarlet* indeed influence *Plodia* testis pigmentation (Heryanto et al. 2022a; Shodja et al. 2026). Additionally, RNA sequencing (RNA-seq) data from *Plodia* gonads across all developmental stages and sexes (Lee et al. 2026) suggests that melanin and ommochrome genes are expressed in *Plodia* larval testis, where phenotypes can be easily observed **(Suppl. Fig. S2)**. To maximize the likelihood of disrupting gene function, we designed sgRNAs primarily against early exons of each target gene, where frameshift mutations are more likely to introduce premature stop codons. An exception was made for genes with multiple transcript isoforms, such as *yellow*, *vermilion*, *kmo/cn*, and *tan*, where sgRNAs were instead designed to target coding regions shared by all isoforms. All selected sgRNA target sites were selected to fall within conserved regions of the corresponding AA sequences (**Figures 1B and 1C)**.

## MELANIN BIOSYNTHESIS PATHWAY

### Ebony and Black regulate pigmentation of larval and pupal integuments and testes

Having identified conserved pigmentation genes and designed corresponding sgRNAs, we first investigated how disruption of melanin pathway genes affects coloration across *Plodia* development. Following early embryonic microinjection of CRISPR/Cas9 components, we examined the pigmentation changes in G_0_ mosaic knockouts (mKOs) at the late larval (L4-L5), pupal, and adult stages and compared them with WT controls. Although *black* and *ebony* affect the same N-β-alanyldopamine (NBAD) branch of the melanin pathway that controls darkening, their disruption led to distinct changes in pigmentation intensity and color hue at all developmental stages (**Figure 2A**). In contrast, *yellow* and *tan* G_0_ mutants exhibited a mild lightening of color that was difficult to distinguish from WT at the larval and pupal stages. Therefore, *yellow* and *tan* G_0_ mutants were only phenotyped only at the adult stage.

**Figure 2.**
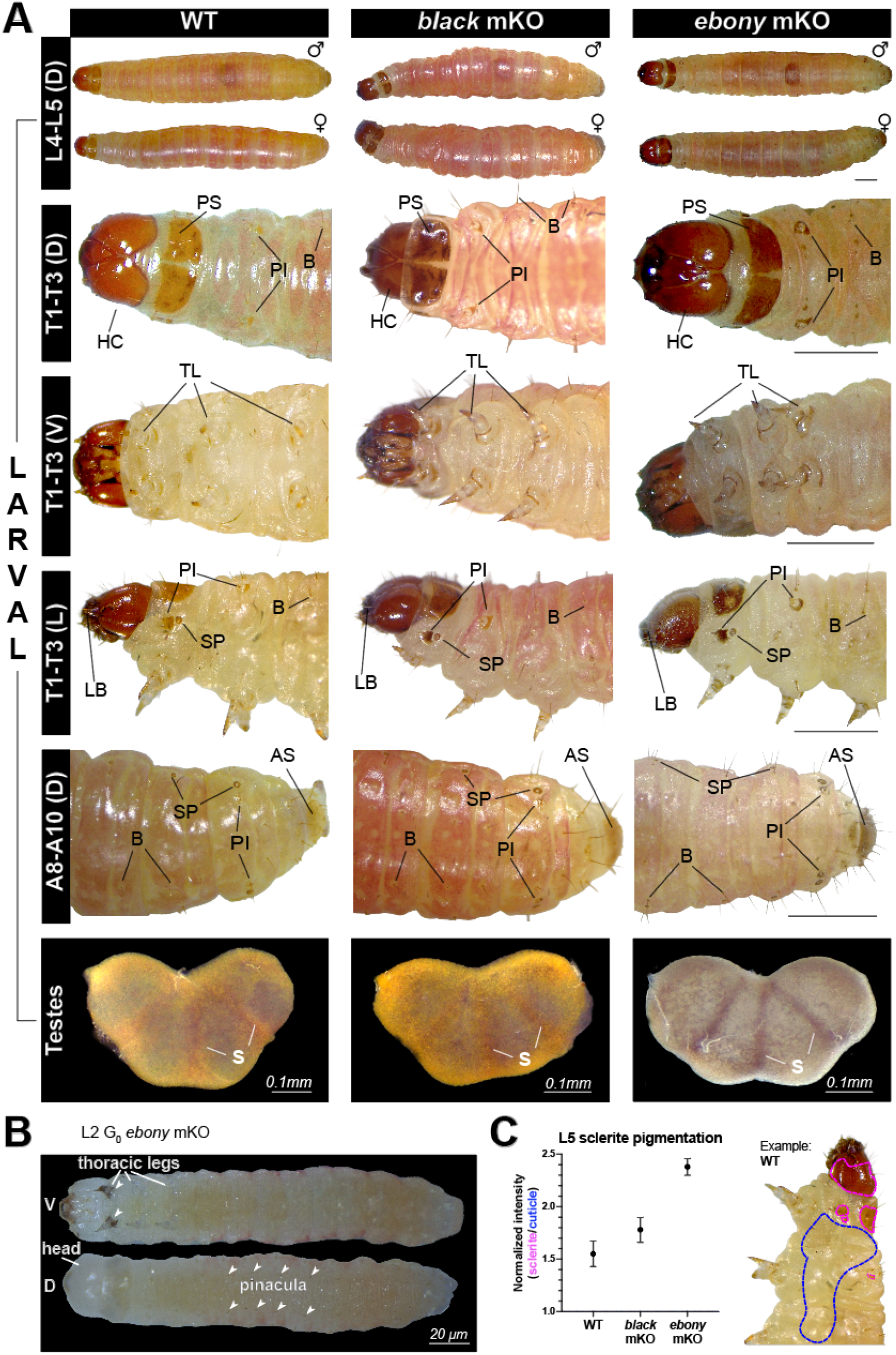
Larval phenotypes resulting from CRISPR/Cas9 knockout of the melanin pathway genes *black* and*ebony*. (A) Phenotypes of *black* and *ebony* mutation on larval cuticles and testes. L4-L5: larval stage 4-5, T1-T3: thoracic segment 1-3, A8-10: abdominal segment 8-10, HC: head capsule, PS: prothoracic shield, PI: pinaculum, B: bristle, TL: true/thoracic leg, LB: labrum, SP: spiracle, AS: anal shield, S: septum, D: dorsal view, V: ventral view, L: lateral view. (B) *ebony* G_0_ KO at the L2 larval stage. The head capsule has no pigment post-molting, but black-tipped thoracic legs are clearly visible against the translucent body. (C) Pigment intensity ratio. Normalized intensity is calculated by dividing the average grayscale of sclerites (magenta) by the non-pigmented region in the larval body (blue) using 1 representative larva image per genotype. Imaging was done in an ambient LED light room. Error bars reflect normalized pigment intensity variation across multiple sclerites regions within an individual.

In larvae, pigment differences between *black* and *ebony* mKO were most obvious in sclerotized structures, while cuticle redness varied with the timing of molting. Although *black* mKO larvae generally displayed a darker red cuticle than WT, *ebony* mKO larvae showed cuticle pigmentation comparable to WT (**Figure 2A**, L4–L5). However, the most striking phenotype observed in both mutants was dark brown pigmentation of the sclerites, such as the prothoracic shield/notum and head capsule (**Figure 2A**, T1–T3), which was visible without magnification (**Suppl. Fig. S3A**). These distinctive phenotypes allowed for the selection of G_0_ *black* and *ebony* mutants as early as the L2 larval stage (**Figure 2B**). Similar pigmentation differences were observed in the thoracic legs, pinacula, spiracles, bristles, and anal shield of larvae (**Figure 2A**), with *black* mKO exhibiting a warmer brown tone and *ebony* mKO displaying a cooler, smoky brown tone of varying intensity.

In addition to these pigmentation changes, *ebony* mKO larvae had visibly darker, straighter, and more rigid bristles than WT (**Figure 2A**, L4–L5). The darkening was most pronounced at the base of the bristles and their surrounding pinacula plate (**Figure 2A**, A8–A10). The labrum, a mouthpart structure that is typically dark brown in WT, appeared almost black in both *black* and *ebony* KO larvae (**Figure 2A**, T1-T3). To quantify these pigmentation differences, we measured overall melanin intensity in the *black* and *ebony* larval mKO using the grayscale values of the prothoracic shield and pinacula normalized against non-pigmented cuticles. Both *black* and *ebony* mKO phenotypes had notably darker pigmentation than WT larvae (**Figure 2C**).

Given the pronounced pigmentation changes in external larval structures, we next examined whether these effects extended to the testes, which are visible through the dorsal cuticle and serve as a distinguishing feature for sex identification in *Plodia* (**Figure 2A**, Testes). Testis pigmentation is also of particular interest as a potential visible indicator of germline CRISPR edits in G_0_ individuals. Because *yellow* and *tan* mutants could not be reliably distinguished from WT during the larval stages, their testes were not examined. However, *ebony* and *black* mKO testes were substantially darker and appeared more opaque through the larval dorsal cuticle than their WT counterpart (**Figure 2A**, Testes). Although *black* mKO testes retained a similar overall tone to WT, their pigmentation was much more saturated, particularly along the three septa separating the four testicular lobes. In contrast, *ebony* mKO testes took on a distinct smoky, gray-brown color.

The pigment changes observed in *black* and *ebony* mKO larvae also extended into pupal development, altering both the timing and appearance of melanization. In WT *Plodia*, pupal development is divided into 14 stages, characterized by progressive pigmentation of the eyes (P1-P9), then the body (P10-P14), before finally eclosing as young adults (Zimowska et al. 1991; Heryanto et al. 2025). Spiracle melanization normally develops alongside the surrounding pupal cuticle and is not readily distinguishable. However, in contrast to WT, both *black* and *ebony* mKO pupae exhibited premature body melanization, characterized by widespread darkening of the dorsal puparium and pronounced pigmentation of the spiracles before eye pigment development was complete (**Figure 3A, 3B, Suppl. Fig. S3B**). This early melanization resembles the phenotype of the *B. mori blackpupa (bp)* mutant, in which disruption of *aspartate decarboxylase* (*BmADC),* the ortholog of *black*, results in melanin deposition as early as three hours after the start of pupation (Dai et al. 2015). By the mid-pupal stage, *black* mKO pupae developed a dark reddish-brown hue with black spiracles (**Figure 3A**). Conversely, the surviving *ebony* mKO pupae displayed a smoked gray-brown color lacking any warm tones, and their puparium appeared wrinkled, rigid, and less plump. Similar phenotypes have been reported after *ebony* KO in the wax moth *G. mellonella* (Luo et al. 2025) and the tea geometrid *Ectropis grisescens* (Li et al. 2024). Together, these findings suggest that loss of either *black* or *ebony* triggers premature pupal melanization, while *ebony* disruption is also associated with altered pupal morphology, potentially reflecting defects in cuticle sclerotization that impact development.

**Figure 3.**
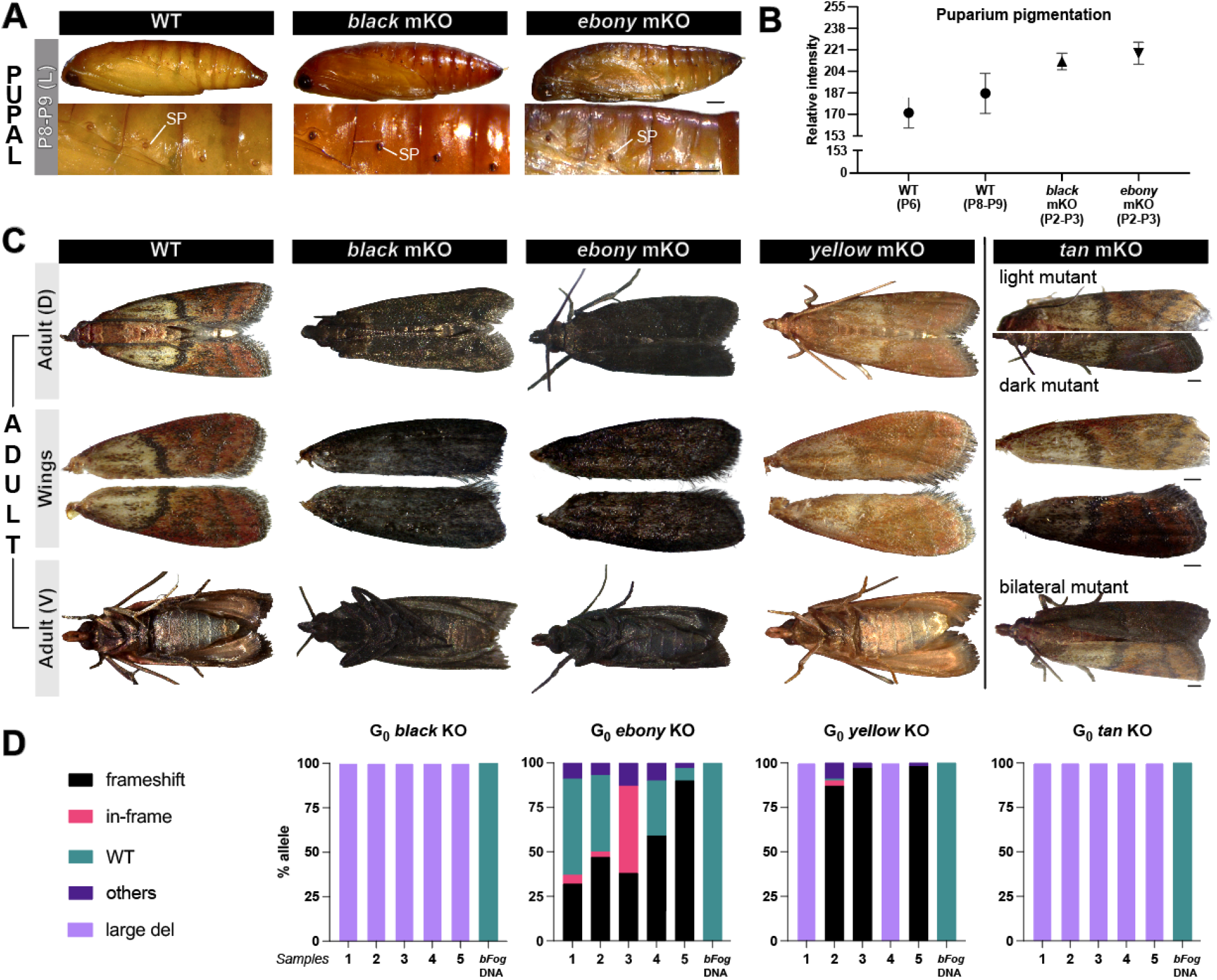
Pupal and adult phenotypes and genotypes resulting from CRISPR/Cas9 knockout of the melanin pathway genes *black*, *ebony*, *yellow*, and *tan*. (A) Phenotypes of *black* and *ebony* mutation at the mid-pupal stage (P6-P9), when eyes are fully pigmented, but the color of the pupal exoskeleton has not developed. SP: spiracle. (B) Puparium pigmentation development of early *black* and *ebony* mKO pupae (P2-P3) in comparison to mid WT pupae (P6-P9). Reference images of these pupae are in **Suppl. Fig. S3B**. (C) Phenotypes of *black*, *ebony, yellow,* and *tan* mutation on adult scales. (D) Parsed alleles in the gonadal tissue of G_0_ crispants. In/del <21 bp were categorized as either frameshift (black) or in-frame (magenta), WT: WT allele (cyan), large del: amplicons too short for alignment or in/dels ≥21 bp (light purple), others: unparsed noise in the sequencing trace (dark purple). Scale bar: 500 µm, unless otherwise noted.

### Black, Ebony, and Yellow differentially regulate adult wing and body pigmentation in Plodia

Although *black* and *ebony* mKOs showed distinct pigment phenotypes at the larval and pupal stages, their adult wing and body coloration was nearly indistinguishable. In both mutants, the scales across the wings and body that are normally beige and reddish in WT (basal/AM, M, and ST regions) were replaced by black and dark gray scales (**Figure 3C; Suppl. Fig. S3C**). However, the transverse wing lines (**Figure 3C**, AM, M, PM, and ST), which are normally black in WT individuals, remained distinguishable as darker bands against the surrounding scales. Loss of *yellow* function showed the opposite phenotype, where black transverse lines lightened to a golden color and beige/red wing regions turned pink. Despite these pronounced changes in saturation and brightness, the overall wing patterning was preserved, and the coloration in the basal/AM area remained unaffected.

Together, these phenotypes are consistent with distinct roles for *black*, *ebony*, and *yellow* in regulating the balance between dark and light melanated pigments in the adult wing scales. The darkening of normally beige/reddish regions after disruption of *black* and *ebony* suggests that these genes promote the conversion of dopamine to NBAD-sclerotin synthesis, limiting dark melanin formation in the wings (Sugumaran and Barek 2016; Zhang et al. 2017a). The persistence of red coloration in the M and ST areas of WT wings (**Suppl. Fig. S3C)** may also reflect contributions from another pigment class, likely the ommochrome pathway. Such spatial differences in pigmentation could involve regulation by similar mechanisms to those mediated by the gene *Optix*, a transcription factor involved in coordinating wing color patterning in butterflies (Zhang et al. 2017b; Banerjee et al. 2024). Conversely, the loss of dark pigment in *yellow* mutants is consistent with a role for Yellow in promoting melanin formation in normally black wing regions. Together, these findings support a model where region-specific melanin pathway gene activity contributes to the distinct color of adult *Plodia* wing scales.

### tan knockout shows a spectrum of pigmentation phenotypes

In contrast to the relatively consistent adult pigment phenotypes observed in *black*, *ebony*, and *yellow* mutants, disruption of *tan* produced a broader spectrum of wing color patterns. Annotation of the *Plodia* genome based on RNA-seq data (Lee et al. 2026) identified three predicted transcript variants of the *tan* gene, arising from three distant transcription start sites (**Suppl. Fig. S2D**, t1, t2, t3). These transcripts encode similarly-sized Tan isoforms, ranging from 390 to 401 AA. Three predicted Tan isoforms were also identified in each of the other lepidopteran species examined suggesting that this alternative splicing pattern may be conserved across the order. However, the functional role of the individual Tan isoforms remains unexplored.

To ensure that all three predicted *tan* transcripts were disrupted, we designed sgRNAs targeting the shared exon 6. Unlike *black* and *ebony* mKOs, tan mKO larvae (L4-L5 G_0_) did not display any distinct pigmentation changes in the sclerites, legs, testes, or cuticles, making the mutant larvae indistinguishable from WT. Thus, the injected G_0_ individuals were allowed to develop into pupae and adults without any phenotypic selection.

In G_0_ adults, the *tan* mKOs displayed a range of pigment phenotypes, including both overall pigment lightening and darkening in the wing scales (**Figure 3C, Suppl. Fig. S3D,** *tan* mKO). Among the 24 surviving individuals, we identified four phenotypic classes. (1) Individuals with a light phenotype (n=13; 6 females, 7 males), which was the most common phenotype in both males and females. These individuals maintained the WT wing patterns and beige scales in the antemedial area, but the overall wing color intensity was slightly reduced. Additionally, the black transverse wing pattern lines (AM, M, PM, and ST) became gray, and red AM and M areas became orange. (2) Individuals with a dark phenotype, which was only observed in males (n=8). This category was characterized by the transverse lines retaining the WT pattern and black color, but the red-scaled area (AM and M) became darker red (maroon) relative to WT. (3) Mixed mosaic phenotype (n=1 male). This individual had gray transverse lines and dark red AM and M areas. Finally, (4) bilateral mosaic phenotype (n=2, one male and one female). These individuals exhibited a sagittal split between dark and light phenotypes. The dark side of the wing turned completely gray and dark brown, losing its beige and black pigments, while the light side of the wing lost its red and black, leaving only beige, gray, and orange pigment. Overall, all *tan* mKO individuals appeared to entirely lose their pigment contrast in the scales, shifting either toward a light, warm tone or a dark, cool tone.

## OMMOCHROME BIOSYNTHESIS PATHWAY

Although disruption of melanin pathway genes produced distinct pigmentation phenotypes across *Plodia* development, their suitability as visible co-CRISPR markers varied with developmental stage and phenotype detectability. To expand the range of potential markers, we next investigated the ommochrome pathway, comparing the effects of disrupting pigment biosynthesis genes (*vermilion* and *kmo/cn*) with those of pigment transport genes (*white* and *scarlet*).

### Ommochrome pathway disruption eliminates red larval pigmentation but produces distinct testis phenotypes

To compare the effects of disrupting ommochrome biosynthesis and transport on larval pigmentation, we generated *vermilion* and *kmo/cn* mKOs, along with previously described *scarlet* and *white* mKOs as controls (Heryanto et al. 2022a; Shodja et al. 2026). At all larval stages, the sclerites and bristles remained unaffected in *vermilion*, *kmo/cn*, *scarlet*, and *white* mKO, suggesting that these genes are not involved in cuticle sclerotization or pigmentation in those tissues. However, all four knockouts resulted in the elimination of red pigments from the larval integument (**Figure 4A**). Similarly, in *kmo/cn* and *scarlet* mKOs, the red gonad pigmentation was reduced to a deep golden color. This remaining pigment in the gonads combined with the loss of cuticle pigment in *kmo/cn* and *scarlet* mKOs at L4-L5 stages still allowed us to visualize the dorsal testicular mass (**Figure 4A**, orange arrows) and ovaries (**Figure 3A**, magenta arrows), enabling sexing based on the shape of these yellow dorsal masses. The two ovaries appeared as separate yellow masses elongated transversely, whereas the testes appeared close together in the center of the A5 segment. Conversely, in *white* mKO mutants, the ovaries and testes lost nearly all pigmentation and appeared translucent under the cuticle (**Figure 4A**, gray arrows), making sexing the larvae extremely challenging. The testes of *vermilion* mKOs appeared pale yellow rather than the colorless phenotype observed in *white* mKOs (**Figure 4A**, Testes). Without *vermilion*, there would be a cytosolic accumulation of L-tryptophan, which, in its crystallized form, is yellowish white (CAS 73-22-3). Pterin, which was reportedly expressed in *Drosophila* testis sheath (Ferré et al. 1986; Ferré 2024), may further contribute to the yellow pigmentation unmasked by the lack of ommochrome in the *vermilion* KO in *Plodia*. *kmo/cn* mKO testes phenocopied that of *scarlet* mKO testes, in which the overall capsule and septa became light yellow, and opacity increased due to the loss of red-brown pigment in comparison to WT (**Figure 4A**, Testes).

**Figure 4.**
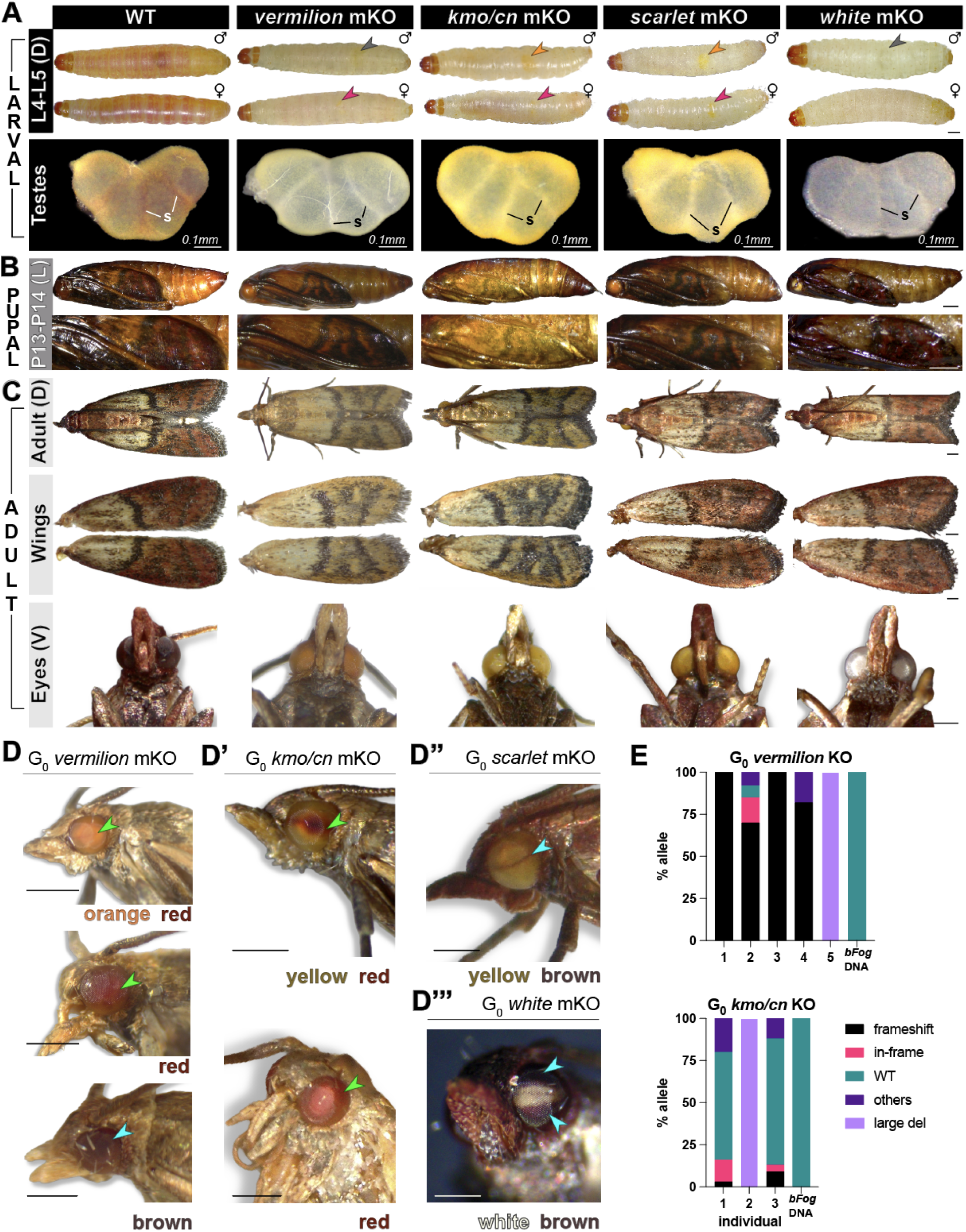
Larval, pupal, and adult phenotypes after CRISPR-mediated knockout of the ommochrome pathway genes *vermilion*, *kmo/cn*, *scarlet*, and *white*. The phenotypes of *vermilion*, *kmo, scarlet,* and *white* mutation (A) on larval cuticles and testes, (B) at pupal stage, and (C) on adult scales and most common eye colors. S: septa; gray arrows: translucent testes, orange arrows: yellow testes, magenta arrows: ovaries. (D) Various eye phenotypes found in G_0_ mosaic KO. Green arrow: red clone, cyan arrow: WT clone. (E) Parsed alleles in the gonadal tissue of G_0_ crispants. In/dels <21 bp were categorized as either frameshift (black) or in-frame (magenta), WT: WT allele (cyan), large del: amplicons too short for alignment or in/dels ≥21 bp (light purple), others: unparsed noise in the sequencing trace (dark purple). Scale bar: 500 µm, unless otherwise noted.

Together, these results indicate that disruption of ommochrome biosynthesis and transport genes produces a common loss of red larval integument pigmentation but distinct testis phenotypes. These differences in gonadal coloration and visibility may be particularly relevant when evaluating pigmentation genes as markers for identifying CRISPR-edited larvae.

### Vermilion and KMO/Cinnabar disruption alters pigmentation in adult wing scales and eyes

Although disruption of *vermilion*, *kmo/cn*, and *scarlet* produced similar larval integument phenotypes (**Figure 4A**), their effects diverged during pupal and adult development. Unlike *white* and *scarlet* mKOs, which visibly altered eye pigment but not wing pigment or morphology, *vermilion* and *kmo/cn* mKO affected both tissues during the pupal and adult stages (**Figure 4B, 4C**). In adult wings, the beige and black regions such as basal/AM and transverse lines remained unchanged (**Suppl. Fig. S3C**). However, red scales were replaced by shades of yellow (**Figure 3C**, wings), resulting in a distinct yellow-and-black wing phenotype. The wings of *vermilion* mutants appeared slightly paler than those of *kmo/cn,* consistent with the downstream role of KMO/Cn in producing yellow-colored 3-OHK. To our knowledge, these findings provide the first genetic evidence that the ommochrome pathway is active in *Plodia* wings, extending the previously established roles of this pathway in eye and testis pigmentation.

The two biosynthesis mutants also exhibited distinct eye pigment phenotypes. Most G_0_ *vermilion* and *kmo/cn* mKOs developed orange and yellow eyes, respectively (**Figure 4C**, eyes) (Shodja et al. 2026). However, a small subset of *vermilion* and *kmo/cn* crispants exhibited red ommatidial patches or entirely red eyes, despite sharing the same yellow-and-black wing phenotypes (**Figure 4D, 4D’**). The appearance of red pigmentation is notable for two reasons: no comparable red stage was observed during normal WT pupal eye development, and red ommatidial patches were not observed in either *white* or *scarlet* mKOs (**Figure 4D’’, 4D’’’**). Similar red eye phenotypes have been described in *kmo/cn* KO of *P. xylostella*, where yellow eyes were associated with frameshift mutation and red eyes were associated with in-frame mutations (Xu et al. 2020).

In contrast, the eye mosaics observed in *scarlet* and *white* mKOs consisted of yellow-and-brown and white-and-brown ommatidia, respectively, without detectable red patches (**Figure 4D″, D‴**). The persistence of brown ommatidia is consistent with incomplete or spatially restricted disruption of ommochrome pigmentation in these G_0_ mosaics. Together, these findings reveal differences between ommochrome biosynthesis and transport mutants in both the affected tissues and the range of eye pigment phenotypes.

## EVALUATION OF PIGMENT GENES AS CO-CRISPR SELECTION MARKERS

### Embryo hatch rates and adult mutant recovery differ among pigmentation gene targets

Although disruption of pigmentation genes produced a range of visible phenotypes across *Plodia* development, their suitability as co-CRISPR markers also depends on the recovery and viability of edited individuals and their ability to transmit mutations to subsequent generations. To evaluate these properties, we first measured embryo hatch rates and adult mutant recovery following CRISPR/Cas9 targeting of these four melanin pathway genes (*black*, *ebony*, *yellow*, and *tan*) and four ommochrome pathway genes (*vermilion*, *kmo/cn*, *white*, and *scarlet*). We recorded embryo hatch rates and the recovery of phenotypically identifiable adult mutants across injection batches. For G_0_ injectees of *black* and *ebony*, individuals were first screened for pigmentation phenotypes at the larval L5 stage and again at the pupal and adult stages. Adult mutant recovery was calculated as a percent relative to the number of L1 hatchlings.

Among melanin pathway targets, *black* injections (674 embryos across three batches) yielded an average hatch rate of 18% (egg to L1) and an adult mutant recovery rate of 23% (L1 to adult; **Table 1**). *ebony* injections (622 embryos, three batches) produced comparable rates of 24% hatching and 25% mutant adult recovery. *yellow* injections (293 embryos, two batches) yielded a 31% hatch rate and a 28% mutant adult recovery. Interestingly, *tan* injections (514 embryos, three batches) produced the highest hatch rate (34%) but adult mutant recovery was the lowest (14%).

**Table 1.** Genome editing results following CRISPR/Cas9 targeting of pigmentation pathway genes.

| Gene targets | Injection time AEL (min) | Total injected (n) | G0 L1 (n) | Hatch rate (%) | Pupal/Adult KO (n) | Mutant adults (%) |
|---|---|---|---|---|---|---|
| *melanin pathway* |  |  |  |  |  |  |
| *black* | 24-51 | 294 | 40 | 14% | 7 | 18% |
|  | 22-52 | 237 | 22 | 9% | 7 | 32% |
|  | 17-49 | 143 | 57 | 40% | 13 | 23% |
| ***black*** |  | **674** | **119** | **18%** | **27** | **23%** |
| *ebony* | 19-50 | 260 | 38 | 15% | 10 | 26% |
|  | 16-45 | 214 | 57 | 27% | 23 | 40% |
|  | 20-54 | 148 | 57 | 39% | 5 | 9% |
| ***ebony*** |  | **622** | **152** | **24%** | **38** | **25%** |
| *yellow* | 30-53 | 168 | 55 | 33% | 15 | 27% |
|  | 22-38 | 125 | 35 | 28% | 10 | 29% |
| ***yellow*** |  | **293** | **90** | **31%** | **25** | **28%** |
| *tan* | 17-40 | 90 | 33 | 37% | 2 | 6% |
|  | 23-52 | 113 | 33 | 29% | 7 | 21% |
|  | 22-57 | 311 | 109 | 35% | 15 | 14% |
| ***tan*** |  | **514** | **175** | **34%** | **24** | **14%** |
| *ommochrome pathway* |  |  |  |  |  |  |
| **vermilion** | **11-46** | **237** | **115** | **49%** | **22** | **19%** |
| *kmo/cn* | 19-36 | 100 | 20 | 20% | 10 | 50% |
|  | 20-40 | 112 | 49 | 44% | 8 | 16% |
| ***kmo/cn*** |  | **212** | **69** | **33%** | **18** | **26%** |
| *white* | 35-80 | 543 | 159 | 29% | 33 | 21% |
|  | 13-43 | 130 | 20 | 15% | 5 | 25% |
| ***white*** |  | **673** | **179** | **27%** | **38** | **21%** |
| *scarlet* | 15-42 | 233 | 66 | 28% | 25 | 38% |
|  | 21-45 | 132 | 83 | 63% | 22 | 27% |
| ***scarlet*** |  | **365** | **149** | **41%** | **47** | **32%** |

Among the ommochrome pathway genes, *vermilion* injections (237 embryos) yielded a 49% hatch rate and 19% adult mutant recovery. *kmo/cn* (212 embryos) and *white* (673 embryos) yielded 33% and 27% hatch rates, and 26% and 21% adult mutant recovery rates, respectively. *scarlet* injections (365 embryos) showed the highest success rates, at 41% hatching and 32% adult mutant recovery. Together, these differences suggest that the recovery of visibly edited adults varies slightly among pigmentation gene targets.

### CRISPR editing generates distinct mutation profiles across pigmentation gene targets in germline tissues

Next, we sought to identify the exact mutations found in the G_0_ individuals and assess whether CRISPR editing occurred in gonadal tissues. Pigmentation phenotypes provide readily observable evidence of somatic editing, but they do not definitively determine the types of mutations present in germ cells and whether they are heritable. To characterize the mutations found in gonadal tissues, we amplified and Sanger-sequenced CRISPR target regions from fused testes or germaria in G_0_ mosaic individuals and analyzed the resulting sequencing data using EditCo ICE v4 (Conant et al. 2022). Mosaic alleles were then categorized by mutation type (in-frame, frameshift, large deletion, etc.) as shown in **Figures 3D** and **4E**. CRISPR targeting of *black* (n=5) and *tan* (n=5) produced mutation profiles that could not reliably be aligned to the reference genome. PCR analysis revealed either smaller-than-expected amplicons or no detectable amplification at the expected WT size (**Suppl. Fig. S4A**). These results are consistent with possible large deletions around the sgRNA target sites, although failed amplification alone cannot confirm their presence. *ebony* G_0_ gonadal genotyping (n=5), on the other hand, revealed a mixture of predicted frameshift and in-frame deletions, as well as some WT alleles. *kmo/cn* G_0_ crispant gonadal genotyping (n=3) suggested predominantly WT alleles or sequencing profiles consistent with large deletions (**Figure 4E**, bottom).

Genotyping of *yellow* G_0_ crispants revealed efficient cutting at the target locus, resulting in either frameshift mutations or large mosaic deletions **(Figure 3D)**. Surprisingly, these frameshift mutations are consistently 2-basepair (bp) deletions with low-to-no mosaicism in the chromatogram near the expected double-strand break (DSB) point, located 3 bp upstream of the Protospacer Adjacent Motif (PAM) sequence. This type of consistent allele output from CRISPR edits suggests possible microhomology-mediated end-joining (MMEJ) repair mechanisms at this site, promoting frequent deletions between identical microhomology loci (Bae et al. 2014; Momose et al. 2018).

### Pigmentation gene disruption reveals gene-specific effects on viability and reproduction

Having characterized the CRISPR-induced mutations in G_0_ individuals, we next investigated whether these alleles could be transmitted to subsequent generations and how their disruption affected development. To this end, we intercrossed G_0_ individuals displaying pronounced mutant phenotypes and examined their G_1_ offspring. Crosses involving *black*, *ebony*, and *yellow* G_0_ individuals produced a mix of wild-type (WT) and mutant offspring, consistent with germline transmission of CRISPR-induced alleles. Although relatively few *black* G_0_ mKO larvae survived to pupation and adulthood, we successfully recovered G_1_ progeny with homozygous frameshift KO (**Suppl. Fig. S4B,** G_1_ *black* KO). These G_1_ mutants displayed L5 larval phenotypes similar to G_0_ crispants (**Figure 2A**). Despite accelerated and extensive melanization at pupal stage, G_1_ mutant puparia remained plump and adults successfully eclosed. In contrast, *ebony* G_1_ mutants exhibited prolonged larval development compared to their WT siblings, and these larvae either died during the L4-L5 stage due to tissue necrosis (**Suppl. Fig. S4C,** left) or at the pupal stage due to failure to complete metamorphosis (**Suppl. Fig. S4C,** right). Sequencing of the G_1_ carcasses revealed biallelic frameshift deletions in *ebony* (**Suppl. Fig. S4B,** G_1_ *ebony* KO).

Germline transmission was also observed following *yellow* disruption. G_0_ *yellow* crispants were intercrossed immediately after eclosion. Although *yellow* mKO males displayed reduced or delayed courtship behavior, these crosses successfully generated G_1_ progeny. Sequencing of G_1_ individuals with *yellow* KO phenotype (n=6) revealed identical genotypes: 2-bp frameshift deletion flanked by microhomologies (**Suppl. Fig. S4B,** blue boxes). As described above, this deletion occurred near short regions of sequence microhomology, consistent with a possible contribution of MMEJ.

In contrast to the other melanin pathway gene targets, *tan* disruption seems to have led to sex-specific phenotypes, albeit with low recovery of adults after injection. Among the *tan* G_0_ crispants, males exhibited a more prominent dark [tan] phenotype and survived to adulthood. In contrast, surviving females showed either subtle lightening or a bilateral mosaic phenotype. Given the Z-linked location of *tan* (**Suppl. Fig. S2**), these differences may reflect sex-dependent effects of gene disruption, although the available data do not establish whether particular *tan* mutations affect viability. To assess transmission, dark phenotype *tan* G_0_ KO males were randomly crossed with their female siblings with subtle phenotypes. The resulting G_1_ progeny displayed predominantly WT pigmentation, and sequencing revealed heterozygous genotypes composed of one WT allele and one 7-bp deletion at the *tan* locus in all four genotyped individuals (**Suppl. Fig. S4B**), suggesting that a WT allele could rescue the *tan* disruption.. As with *yellow*, short regions of microhomology flanked the deletion site (**Suppl. Fig. S4B,** blue boxes), suggesting that MMEJ may have contributed to its formation. However, because only a subset of G_0_ and G_1_ individuals were sequenced, the full diversity and inheritance patterns of *tan* alleles remain unresolved.

Finally, intercrosses between *kmo/cn* G_0_ mutants produced G_1_ progeny displaying either a WT or a pronounced KO phenotype (yellow eyes and yellow-and-black scales). No red-eyed phenotype was observed in any G_1_. Unexpectedly, sequencing of all four examined full-KO-phenotype G_1_ individuals revealed the presence of both WT and frameshift alleles (**Suppl. Fig. S4B**), rather than the biallelic frameshift genotype expected for a true homozygous null. Additional genotyping across subsequent generations will be necessary to resolve the relationship between *kmo/cn* genotype, pigmentation phenotype, and viability.

### white and scarlet serve as effective co-CRISPR markers for yellow disruption

Among the eight pigmentation genes examined in this study, *scarlet* and *white* emerged as the top target candidates for co-CRISPR markers because their disruption produced easily identifiable phenotypes, relatively high recovery rates of visibly mutant adults, and limited apparent effects on development and reproduction. To evaluate their ability to predict editing at a second locus, (**Figure 5A**) we selected the *yellow* gene as a proxy GOI, since *yellow* disruption has visible, scorable, non-lethal effects and no phenotypic overlap with *white* or *scarlet* disruption. This combination allowed us to assess the co-occurrence of mutant phenotypes in G_0_ adults without sequencing every survivor.

**Figure 5.**
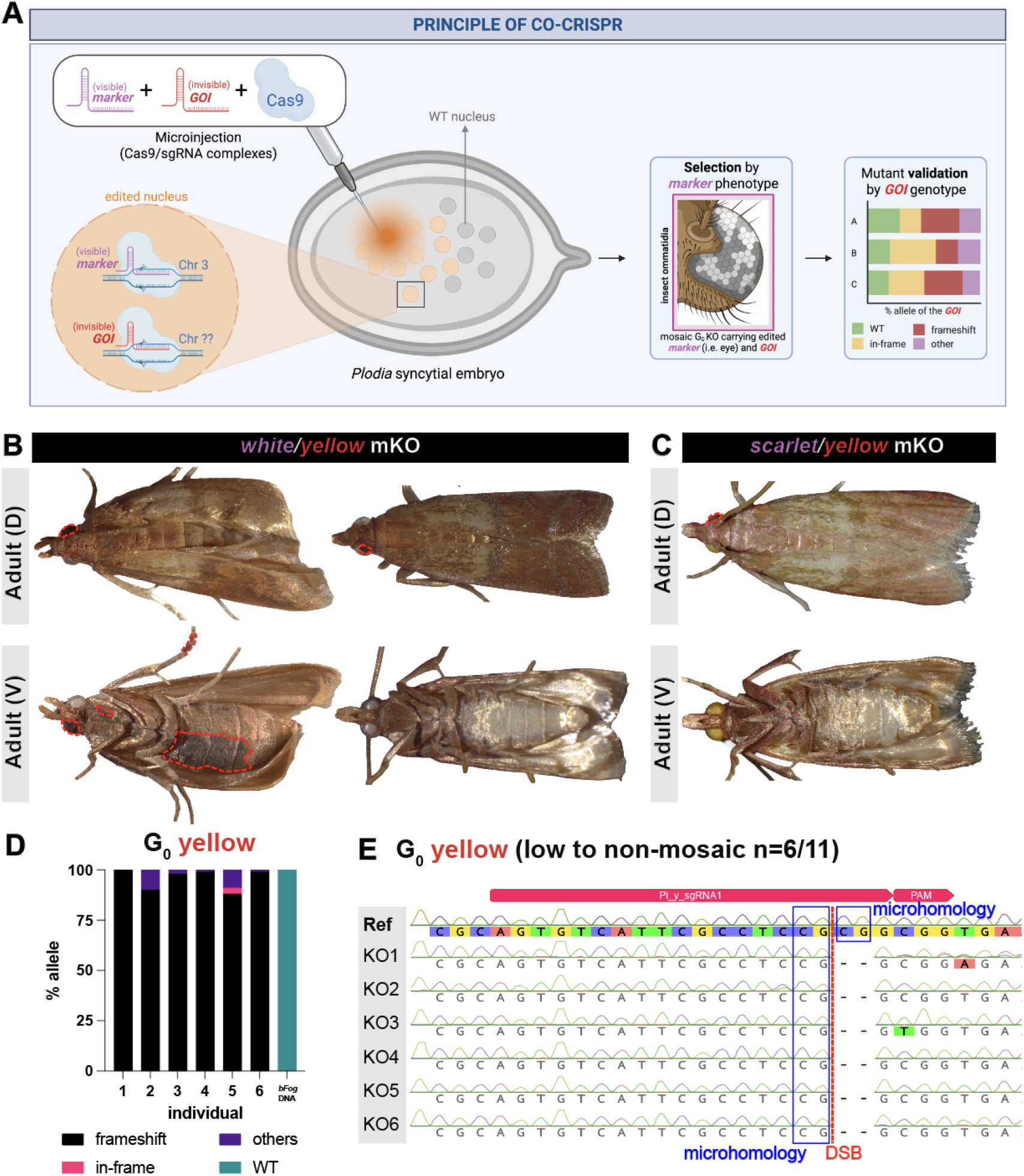
Co-CRISPR validation and Sanger sequencing are consistent with potential microhomology-mediated end joining (MMEJ) repair in *yellow* knockouts. (A) Principle of co-CRISPR in *Plodia* syncytial embryo. When nuclei are targeted during syncytial blastoderm stage with Cas9/marker and Cas9/GOI complexes, a subset of nuclei are likely to be edited at both loci within the same individual. Marker gene is used as an enrichment tool during selection. The GOI edits are validated by sequencing. The schematic was created using Biorender. (B) KO phenotype of co-CRISPR of *white/yellow.* Red dotted lines show WT clones on the eye and body (left) and eye (right). (C) KO phenotype of co-CRISPR of *scarlet/yellow.* Red dotted lines show WT clones. (D) *yellow* G_0_ KO genotype of gonadal tissue. (E) G_0_ *yellow* sequencing chromatograms. Microhomologies (blue boxes) found in *yellow* target flank low to non-mosaic 2-nt cuts in the proximity of DSB (red dashed line). GOI: gene of interest, D: dorsal view, V: ventral view, MMEJ: microhomology-mediated end joining, PAM: Protospacer-Adjacent Motif, DSB: double-strand break.

We first co-injected sgRNAs targeting *white* and *yellow* at equal concentrations. The resulting embryo hatch rate (32%, **Table 2**) was comparable to *yellow* sgRNA alone (31%, **Table 1**) and higher than that of *white* sgRNA alone (27%, **Table 1**). To assess the predictive power of *white* as a co-CRISPR marker, we scored eye and body pigmentation in surviving G_0_ adults and classified individuals using a 2x2 contingency table (**Table 3**). Individuals with WT eyes and WT body were classified as true negatives, whereas those displaying both [white; brackets refers to phenotype of the corresponding mutant gene] eyes and [yellow] body mutant phenotypes were classified as true positives. Individuals with [white] eyes but WT body pigmentation were classified as false positives, and those with WT eyes but [yellow] bodies were classified as false negatives.

**Table 2.** CRISPR mutagenesis outcomes of wild-type *Plodia interpunctella* eggs co-injected with *white* or *scarlet* (markers) and *yellow* (proxy GOI) sgRNA.

| Co-injected gene targets | Injection time AEL (min) | Total injected (n) | G<sub>0</sub> L1 (n) | Hatch rate (%) | Pupal/Adult (n) | eye G<sub>0</sub> mutants | *yellow* G<sub>0</sub> mutants |
|---|---|---|---|---|---|---|---|
| *white + yellow* | 11-37 | 495 | 87 | 18% | 40 | 26 (65%) | 100 (57%) |
|  | 14-40 | 492 | 209 | 42% | 135 | 81 (60%) |  |
|  | 20-50 | 437 | 165 | 38% | 117 | 49 (42%) | 68 (58%) |
| ***white + yellow*** |  | **1424** | **461** | **32%** | **292** | **156 (53%)** | **168 (58%)** |
| *scarlet + yellow* | 15-48 | 410 | 74 | 18% | 34 | 8 (24%) | 12 (35%) |
|  | 20-66 | 349 | 94 | 27% | 27 | 8 (30%) | 8 (30%) |
| ***scarlet + yellow*** |  | **759** | **168** | **22%** | **60** | **16 (27%)** | **20 (33%)** |

**Table 3.** Phenotype (2x2) scoring matrix and analysis of *white+yellow* and *scarlet+yellow* co-CRISPR in adult *Plodia interpunctella*.

| **Phenotypes** |  | **Marker unedited (WT)** |  |  | **Marker edited** |  |  |
|---|---|---|---|---|---|---|---|
| **GOI unedited (WT)** |  | True negative (TN) |  |  | False positive (FP) |  |  |
| **GOI edited [yellow]** |  | False negative (FN) |  |  | True positive (TP) |  |  |
| *Injection -->* |  | *white* + *yellow* sgRNA |  |  | *scarlet* + *yellow* sgRNA |  |  |
| **Phenotypes** |  | **Eyes** |  | *Total* | **Eyes** |  | *Total* |
|  |  | **WT** | **[white]** |  | **WT** | **[scarlet]** |  |
| **Body** | **WT** | **35** | *0 | 35 | **22** | *0 | 22 |
|  | **[yellow]** | **19 | **81** | 100 | **4 | **8** | 12 |
| *Total* |  | 54 | 81 | 135 | 26 | 8 | 34 |
| *FP %* |  | *0/35 = 0%* |  |  | *0/22 = 0%* |  |  |
| *FN %* |  | *19/(19+81) = 19%* |  |  | *4/(4+8) = 33%* |  |  |
| *PPV %* |  | *81/81 = 100%* |  |  | *8/8 = 100%* |  |  |
| *NPV %* |  | *35/54 = 65%* |  |  | *22/26 = 85%* |  |  |

PPV: positive predictive value (see Materials and Methods).
NPV: negative predictive value (see Materials and Methods).
Adapted from a 2x2 diagnostic table (Altman and Bland 1994; Trevethan 2017).

Most strikingly, the *white* co-CRISPR marker achieved a 100% positive predictive value (PPV) and 0% false positive rate. Out of the 135 total phenotyped G_0_ mutants, all 81 that displayed a [white] eye phenotype also displayed a [yellow] body phenotype, regardless of the apparent size of the mutant eye clone (**Figure 5B**, left; **Table 3**). Thus, selection based on the visible [white] phenotype consistently enriched for individuals exhibiting a second pigmentation mutation.

However, 19 individuals displayed a *yellow* mutant body phenotype without a detectable *white* mutant eye phenotype, indicating that selection based on [white] alone would exclude some individuals with visible edits at the secondary locus. One possible explanation for this discrepancy is differences in editing efficiency or repair/mutation outcomes between the two targets. Alternatively, since mutant G_0_ individuals are likely mosaics, small *white* mutant clones may have escaped detection during scoring, particularly given the smaller area available for scoring eye pigmentation compared with body pigmentation. The definition of the other metrics are further explained in the Materials and Methods section.

We next evaluated *scarlet* as a co-CRISPR marker. When targeted alone, *scarlet* sgRNA injection yielded a high L1 hatch rate (41%) and high adult mutant recovery rate (32%, **Table 1**). Surprisingly, *scarlet+yellow* co-injection yielded a much lower hatch rate (16%, **Table 2**, **Figure 5C**). Despite this lower hatching rate, *scarlet* offers practical advantages as a marker candidate. In particular, *scarlet* disruption retains sufficient gonadal pigmentation to allow visual identification of larval testes and ovaries, whereas *white* disruption produces nearly translucent gonadal tissues that are more difficult to distinguish. Previous work has also identified *scarlet* as a potentially less deleterious pigmentation target than *white* in some Lepidoptera (Shodja et al. 2026).

Despite differences in hatch rates and larval phenotypes, *scarlet* and *white* performed similarly as predictors of visible *yellow* disruption. The proportions of correctly classified G_0_ adults were 86% for *white* and 88% for *scarlet*, with false-negative classifications accounting for 19% and 33% of marker-negative individuals, respectively. Neither marker produced detectable false-positive classifications, yielding a PPV of 100% in both co-injection experiments.

Together, these results indicate that visible disruption of either *white* or *scarlet* provides a reliable means of enrichment for G_0_ individuals with a concurrent *yellow* mutant phenotype, although marker-negative individuals may also carry detectable secondary edits.

### Recurrent yellow deletions in G_0_ crispants and G_1_ progeny are consistent with microhomology-mediated repair

To validate the presence of *yellow* mutations, we extended our analysis of the *white+yellow* co-injection experiment. Sequencing revealed that most of the G_0_ crispants displaying the [white] phenotype also carried frameshift alleles at the *yellow* locus (**Figure 5D**). Notably, a recurrent 2-bp deletion was detected in half of the sequenced G_0_ individuals sequenced (3/6), accompanied by relatively little evidence of mosaicism in the Sanger chromatograms (**Figure 5E**). The recurrence of this deletion near short regions of sequence microhomology suggests that MMEJ repair may have contributed to its formation. Together, these results provide proof of concept that the [white] phenotype can serve as a visible indicator of CRISPR activity and enrich for individuals carrying mutations at a second, non-lethal GOI, such as *yellow*.

In a third batch of *white*+*yellow* co-injections (**Table 2**), G_0_ mutant adults could not be reliably scored for the [yellow] phenotype because routine activities, including courtship, mating, and movement within their rearing containers, resulted in progressive loss of wing scales. Of four intercrosses, each involving two to three pairs of G_0_ mutant adults, only one produced G_1_ progeny. We also observed reduced or delayed courtship behavior in G_0_ males, including limited pursuit of females with wing-flapping behavior, even when nearby females displayed calling behavior characterized by a curved abdomen. Similar behavior has previously been observed in G_0_ *yellow* mutant intercrosses.

To further verify the presence and inheritance of *yellow* mutations, we sequenced the target region of the *yellow* gene of G_1_ offspring from the successful intercross. Sanger chromatogram showed overlapping sequence signals near the expected CRISPR cut site, consistent with the presence of insertion/deletion (in/del) alleles. Chromatogram deconvolution identified two frameshift alleles, including a 2-bp deletion immediately upstream of the sgRNA PAM sequence (**Suppl. Fig. S4B**). Both alleles showed signatures of possible MMEJ repair and were predicted to disrupt Yellow protein function. All 80 G_1_ progeny from this intercross displayed both [yellow] body and [white] eye phenotypes, providing evidence that mutant alleles at both loci were transmitted through the G_0_ germline. Although the severity of somatic mosaic mutant phenotypes in G_0_ individuals does not necessarily predict the genotype of their gametes, these findings further support the utility of *white* as a co-CRISPR marker for selecting G_0_ individuals with an increased likelihood of carrying heritable mutations at a secondary target locus.

### Marker-based co-CRISPR selection enriches for mutations in genes without readily visible phenotypes

To further evaluate the utility of marker-based co-CRISPR selection for genes without readily observable mutant phenotypes, we selected six meiosis-related genes (*SPO11*, *SIX6OS1*, *SYCP3*, *HEI10*, *DMC1*, and *REC8*) (Xiang et al. 2023; Benner et al. 2026) as GOIs and co-targeted each of them with the eye marker gene *white* (**Table 4; Supp. File**). These genes encode proteins involved in key meiotic processes, including DNA double-strand break formation and repair, chromosome pairing and synapsis, crossover formation, and sister chromatid cohesion. Because disruption of these processes may affect fertility without producing obvious external phenotypes, these genes provide useful targets for evaluating co-CRISPR selection.

**Table 4.**
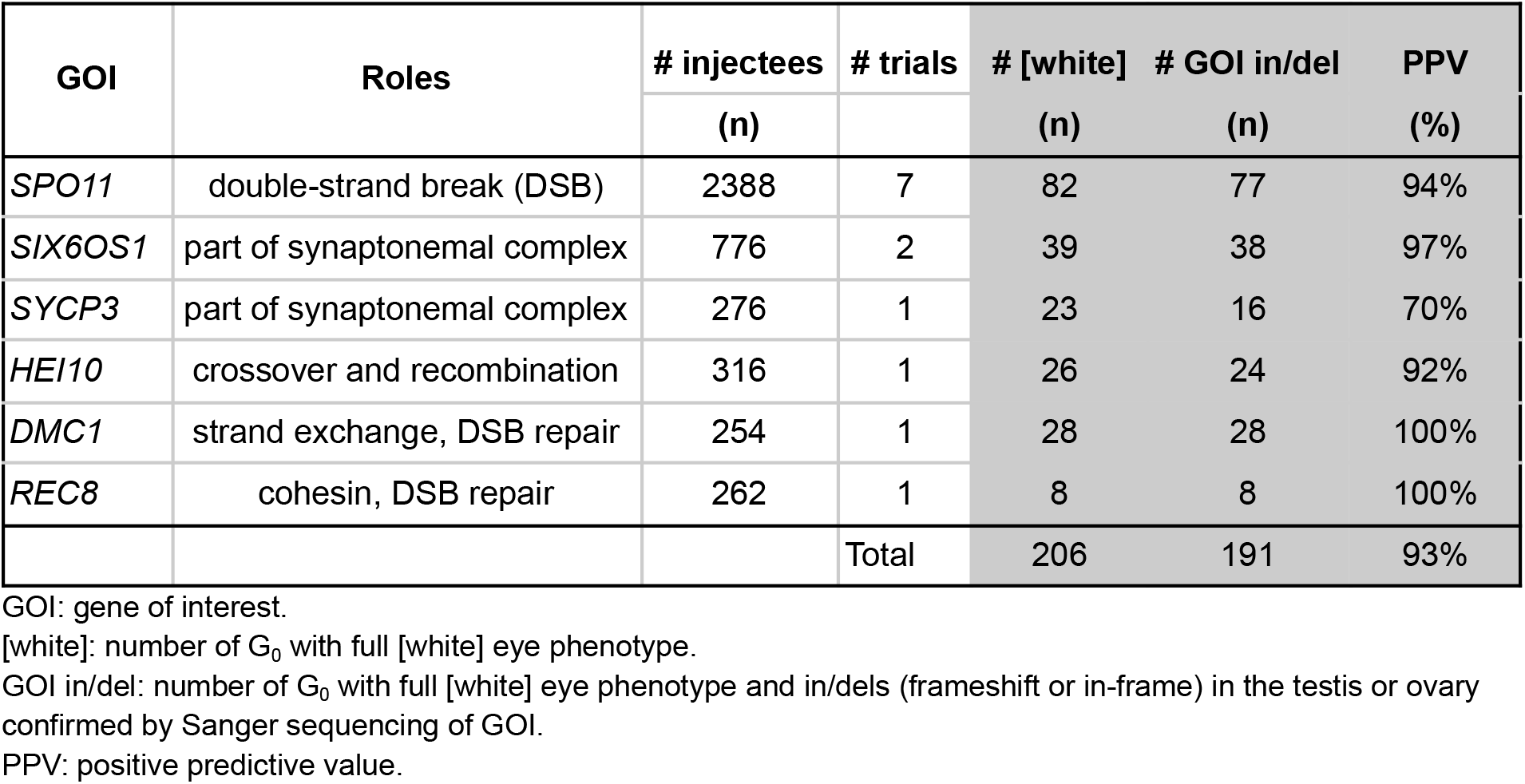
Genotype validation of CRISPR/Cas9 editing in meiosis-related genes using the eye marker *white* for co-CRISPR selection.

| GOI | Roles | # injectees | # trials | # [white] | # GOI in/del | PPV |
|---|---|---|---|---|---|---|
|  |  | (n) |  | (n) | (n) | (%) |
| *SPO11* | double-strand break (DSB) | 2388 | 7 | 82 | 77 | 94% |
| *SIX6OS1* | part of synaptonemal complex | 776 | 2 | 39 | 38 | 97% |
| *SYCP3* | part of synaptonemal complex | 276 | 1 | 23 | 16 | 70% |
| *HEI10* | crossover and recombination | 316 | 1 | 26 | 24 | 92% |
| *DMC1* | strand exchange, DSB repair | 254 | 1 | 28 | 28 | 100% |
| *REC8* | cohesin, DSB repair | 262 | 1 | 8 | 8 | 100% |
|  |  |  | Total | 206 | 191 | 93% |

GOI: gene of interest.
[white]: number of G<sub>0</sub> with full [white] eye phenotype.
GOI in/del: number of G<sub>0</sub> with full [white] eye phenotype and in/dels (frameshift or in-frame) in the testis or ovary confirmed by Sanger sequencing of GOI.
PPV: positive predictive value.

To assess the effectiveness of this approach, we sequenced the target loci in G_0_ individuals displaying the [white] eye phenotype. Selection based on the [white] phenotype successfully enriched for G_0_ individuals carrying sequence-confirmed in/dels at the GOI with PPV values ranging from 70 to 100%. Across the six meiosis-related genes, 191 of 206 [white] G_0_ individuals (93%) carried a confirmed in/del at the corresponding GOI. Five of the six targets yielded PPVs of 92-100%, while *SYCP3* was an outlier at only 70%. This lower PPV may reflect differences in sgRNA cutting efficiency, mosaicism, or other target-dependent factors rather than a general limitation of the co-CRISPR approach, although these possibilities remain to be tested.

## DISCUSSION

In this study, we demonstrated that visible pigment phenotypes provide an effective means of enriching for CRISPR mutants at loci without readily scorable external phenotypes. These pigmentation genes, long used as visible markers in *Drosophila,* function similarly in *Plodia*, producing broadly comparable pigmentation phenotypes (**Suppl. Table S5**). Their effects on fitness, however, varied considerably among genes, ranging from fully viable and fertile KO to severe developmental defects and reduced survival, emphasizing the need to consider potential pleiotropic effects when choosing a marker. Beyond validating these markers, our study provides evidence that the ommochrome biosynthesis pathway contributes to wing pigmentation in *Plodia*.

Among these pigmentation genes, we found that co-CRISPR with *white* or *scarlet* can predict, with exceptionally high PPV (100%), whether a given individual also carries a mutation at an unlinked secondary GOI. Furthermore, selection based on the [white] phenotype identified sequence-confirmed mutations at secondary targets in 93% of G_0_ individuals tested across six meiosis-related genes, offering a practical enrichment strategy for genome editing in *Plodia*. In addition, we identified recurrent deletions consistent with possible MMEJ,although the involvement of this repair pathway requires further investigation using additional targets and larger sample sizes.

### Melanin biosynthesis pathway

Sclerotization and melanization are central to insect survival, and their disruption can have consequences beyond coloration. Sclerotization, or cuticle hardening, is particularly important because insects rely on their exoskeleton for structural support. Immediately after larval molting, the new cuticle is soft and pale. NBAD- and NADA-derived quinones subsequently participate in crosslinking cuticular proteins and chitin fibers (Andersen 2010), converting the pliable cuticle into a rigid exoskeleton that supports mobility and protects the developing adult during pupation in holometabolous insects (Sugumaran and Lipke 1982; Klowden 2013). In adults, these melanin-associated proteins also contribute to wing and body coloration, which can play important roles in survival and reproduction. Pigment patterns may serve as camouflage against predators (Stevens and Merilaita 2008), warning signals that deter attacks (Ruxton et al. 2019), or visual cues involved in mate recognition and courtship (Finkbeiner et al. 2014). Thus, the balance between melanization and sclerotization is important not only for structural integrity, but also for ecological and behavioral functions throughout the insect life cycle (Wittkopp and Beldade 2009).

The larval phenotypes of *black* and *ebony* mKO were most pronounced in sclerotized structures, including the prothoracic shield, head capsule, anal shield, pinacula, and spiracles. Disruption of this pathway could therefore have consequences beyond pigmentation, particularly in structures where proper cuticle hardening is essential. Although we characterized the pigmentation changes, fitness, and fertility associated with loss of function of these genes, several mechanistic questions remain. For example, does altered melanization affect the structural or physiological function of bristles, which may act as mechanosensory receptors involved in feeding and locomotion, or spiracles, which connect the respiratory system to the external environment? Could altered pigmentation of the testis affect germline development and meiotic processes? Finally, might any of these changes contribute to the prolonged larval development observed in some mutants?

Scale color patterning in insects is thought to be regulated in part by the opposing activities of Yellow and Ebony (True et al. 2005; Futahashi et al. 2022), which promote and suppress melanin formation, respectively. Because Tan reverses the Ebony-catalyzed reaction, and given the relatively subtle phenotypes observed in some *tan* mutants, we propose that it helps fine-tune the balance between melanin and NBAD-derived sclerotin rather than acting as an independent regulatory switch.

The diversity of *tan* KO phenotypes observed in our study contrasts with the predominantly lighter color expected following disruption of *tan,* which normally hydrolyzes NBAD into dopamine and β-alanine, thereby antagonizing *ebony* activity (**Figure 1A**). (Wittkopp et al. 2009; Rogers et al. 2014; Camino et al. 2015; Liu et al. 2019; Kuwalekar et al. 2020; Futahashi et al. 2022). The resulting balance between Ebony and Tan enzymatic activity, rather than the action of a single gene alone, may therefore influence melanin and sclerotin production. The broad spectrum of phenotypes may also reflect the three predicted *tan* transcript variants targeted at the shared exon 6 in our experimental design, together with the inherent mosaicism of G_0_ editing. Differences in the extent of editing among cells, potentially combined with transcript-specific effects, could alter the balance between NBAD and dopamine availability, shifting pigment deposition toward darker melanin or lighter NBAD-derived sclerotin in the wings. However, the contributions of individual transcript variants and the underlying metabolic changes remain to be determined.

The sex-associated distribution of *tan* mutant phenotypes may also be related to its Z-linked location in *Plodia*. Pronounced dark pigmentation was observed only among surviving G_0_ males, whereas surviving females exhibited lighter or bilateral mosaic phenotypes. These differences may reflect sex-dependent effects of *tan* disruption, although the relative contributions of gene dosage, mosaicism, and potential effects on viability remain unclear.

Beyond their roles in pigmentation, these melanin-associated genes are also involved in neurophysiological function and behavior, which may contribute to the fitness costs observed in our mutants. In *D. melanogaster*, *tan* and *ebony* play opposing roles in neurotransmitter metabolism at the photoreceptor synapses (Borycz et al. 2002). Ebony converts histamine into carcinine, whereas Tan hydrolyzes carcinine back into histamine for recycling. Given these additional functions, disruption of *tan* may have contributed to the reduced survival observed between larval and pupal or adult stages even among some male G_0_ crispants. *D. melanogaster yellow* mutants also exhibit reduced mating success associated with impaired male courtship behavior and lack of sex comb (Massey et al. 2019). In addition, melanin-associated pathways are involved in immune function through the phenoloxidase cascade, which contributes to wound healing and pathogen encapsulation (Binggeli et al. 2014; Dudzic et al. 2015; Gonzalez-Baez et al. 2025). In our experiments, *tan* disruption was associated with lower adult mutant recovery than *yellow* or *ebony* disruption. Whether this difference reflects reduced viability, differences in editing efficiency, or other factors remains unclear.

### Ommochrome biosynthesis pathway

Across Lepidoptera and other insect orders, *kmo/cn* knockouts have consistently been shown to alter eye pigmentation, producing yellow, red, or otherwise depigmented ommatidia. In the diamondback moth *Plutella xylostella*, for example, *kmo/cn* disruption produced yellow or red eyes depending on the type of mutation (Xu et al. 2020). The availability of circulating pigment precursors may also influence these phenotypes through cell-nonautonomous rescue. In larva of the flour moth *Ephestia kühniella*, an immediate relative of *Plodia*, *kmo/cn* activity has been restricted to the Malphigian tubules, contributing to the production and circulation of ommochrome precursors in the hemolymph (Stratakis 1982). During the pupal stage, ommatidia can take up injected 3-OHK from the hemolymph and use it for ommochrome production (Muth 1969). Such uptake provides a plausible mechanism by which mutant eye cells could acquire pigment precursors synthesized elsewhere.

The similar testicular pigmentation phenotypes of *kmo/cn* and *scarlet* mutants are consistent with their respective roles in 3-hydroxykynurenine (3-OHK) synthesis and transport, both of which contribute to downstream ommochrome pigment production. The greater translucency previously observed in *white* mKO testes may reflect White’s broader role as a partner of multiple ABC transporters, including Scarlet in ommochrome transport and Oily kinshiryu (Ok) in uric acid transport (Zhang HaoKun et al. 2018; Shodja et al. 2026).

Wing pigmentation, by contrast, shows a more variable dependence on the ommochrome pathway across species. In *Plodia*, disruption of *white* (Heryanto et al. 2022a; Shirk et al. 2023) or *scarlet* (Shodja et al. 2026) have been shown to affect L5 cuticle pigmentation, testicular pigmentation, and adult eye coloration, but does not produce comparable changes in wing pigmentation (**Figure 4**). This resembles the tissue-specific differences in ommochrome pathway requirements previously reported in *Bicyclus anynana* (How et al. 2023), where disruption of *vermilion* and *cinnabar* eliminated eye pigmentation but left wing coloration unaltered, despite the presence of ommochromes and their precursors in wing scales. Other species also show differences between eye and wing pigmentation. In the silkworm moth *Bombyx mori*, for example, a mutant affecting an ommochrome-associated cytosolic transporter alters wing and egg coloration but not the eye pigmentation (Uchiyama et al. 2023).

Our findings on Vermilion and KMO/Cn suggest that normal wing pigmentation in *Plodia* depends on the canonical ommochrome pathway, at least through the enzymatic steps leading to 3-OHK production. The retention of normal wing pigmentation following *white* or *scarlet* disruption, however, suggests that subsequent pigment processing or deposition may occur independently of the canonical White-Scarlet transport system. This could involve an alternative transport mechanism or pigment synthesis occurring outside the conventional pigment granules. Consistent with the possibility of an alternative transport system, tryptophan and 3-OHK have been reported to move from the hemolymph into wing cells in *Heliconius* (Reed et al. 2008), while *scarlet* and *white* expression was not detected in red wing regions (Ferguson and Jiggins 2009). Differences may also extend beyond transport. In the nymphalid butterfly *Aglais io*, *cardinal*, which encodes an enzyme involved in the later steps of ommochrome synthesis, was expressed at much lower levels in wings than in eyes (Uchiyama et al. 2023). Together, these observations highlight tissue-specific differences in ommochrome synthesis and transport, raising questions about the requirement for White and Scarlet transporters and the role of pigment granules in wing pigmentation.

### Practical approaches in using co-CRISPR markers

The identification of pigmentation genes suitable for co-CRISPR selection represents a valuable addition to the *Plodia* genetic toolkit, as visible markers greatly simplify the screening of edited individuals. This is especially important because embryonic microinjection in *Plodia* typically yields only tens of survivors from several hundred injected eggs, making rapid visual identification of mutants preferable over molecular genotyping of every surviving individual. In insects, candidate markers are often selected from the melanin and ommochrome pathways, which are well characterized and can produce robust, visually distinct phenotypes upon disruption. An ideal co-target gene should preferably (1) produce a visible and easily scorable phenotype, (2) be viable in the homozygous mutant state, (3) be efficiently disrupted by CRISPR/Cas9 targeting, and (4) have minimal pleiotropic effects that could interfere with the function of the GOI or its associated pathway.

In this study, we designed sgRNAs using established guidelines and tools such as CRISPOR (Concordet and Haeussler 2018). We selected target regions based on predicted transcript variants and sequence conservation across multiple insect species, aiming to disrupt conserved regions of each gene. When available, RNA-seq data can also help identify transcripts expressed in relevant tissues. Targeting multiple transcript variants may produce a broader range of phenotypes, as observed in *tan, vermilion*, and *kmo/cn*. Thus, targeting one variant at a time may be preferable for studying their specific functions, whereas targeting shared exons or using multiple sgRNAs may be more appropriate when broader gene disruption is desired.

The ommochrome genes *white* and *scarlet* meet the criteria established for an ideal co-CRISPR marker, although *scarlet* KO testes and ovaries are easier to identify visually in L5 larvae than *white* KO gonads. To some extent, the marker phenotype may also serve as an indicator of CRISPR editing activity and mosaicism in G_0_ individuals. However, its predictive value depends partly on the relative editing efficiencies of the marker and GOI targets. Although the severity of a G_0_ marker phenotype may not consistently reflect the extent of editing at the GOI, our co-injection experiments demonstrate that selection based on [white] or [scarlet] phenotypes reliably identified individuals displaying concurrent [yellow] phenotypes, with a PPV of 100% for both markers.

Importantly, this approach was also effective for genes without readily scorable external phenotypes. Across six meiosis-related GOIs, 191 of 206 [white] G_0_ individuals (93%) carried sequence-confirmed indels at the corresponding target locus. These results demonstrate the practical value of co-CRISPR selection for identifying G_0_ individuals carrying secondary mutations without requiring molecular screening of every injected survivor. However, the predictive performance varied among GOIs, emphasizing the importance of target-specific sgRNA efficiency and other factors influencing editing outcomes.

After co-CRISPR marker selection, sequencing G_0_ gonadal tissue can provide additional evidence of editing at the GOI. Combined with Sanger chromatogram deconvolution tools such as the web-based EditCo ICE v4 (Conant et al. 2022), this workflow is particularly valuable when studying genes associated with lethality or sterility, including germline and meiosis-related genes, for which generating G_1_ mutants or establishing homozygous stocks may be difficult. However, gonadal sequencing cannot substitute for direct G_1_ genotyping because G_0_ gonadal samples may contain both somatic and germline cells, and mosaic individuals may carry edited alleles in only a subset of their germ cells. Thus, G_0_ gonadal sequencing should be considered an additional validation step for GOI editing rather than definitive evidence of germline transmission.

Another notable observation was the relatively low apparent mosaicism in the some G_0_ *yellow* and *tan* sequencing results, together with recurrent deletions at target sites flanked by short regions of microhomology near the expected double-strand break. These sequence features are consistent with a possible contribution of MMEJ activity during CRISPR-induced DNA repair in *Plodia* embryos (Nakade et al. 2014; Romanowski et al. 2026), This possibility could inform future sgRNA design strategies targeting regions with nearby microhomologies to favor more predictable deletions. It may also motivate evaluation of MMEJ-based knock-in approaches, such as Precise Integration into Target Chromosome (CRISPR PITCh) (Sakuma et al. 2016), in *Plodia*.

Finally, another potential advantage of co-targeting a well-characterized marker alongside the GOI is its use as a reference for evaluating development and survival. For example, the routine recovery of *white* KO mutants in our lab provides a useful benchmark for comparing survival outcomes following co-targeting of secondary genes, particularly those suspected of affecting development or viability. Such comparisons may help identify developmental stages at which a GOI is required, although reduced survival cannot be attributed to GOI disruption without appropriate controls. In practice, CRISPR-associated lethality may also be influenced by sgRNA concentration, injection timing, and embryo survival. Higher sgRNA concentrations or earlier injection times may reduce survival rates through injection-associated toxicity, increased editing activity, or disruption of a gene required for early development. Therefore, a visible co-CRISPR marker with well-characterized survival outcomes may not only be useful for selecting G_0_ individuals but also as a reference when comparing survival rates across experiments.

### Limitations of the study

Several limitations should be considered when interpreting these results. First, we did not comprehensively assess the pleiotropic effects of the pigmentation genes examined in this study beyond coloration and fitness at G_0_. Many melanin and ommochrome pathway genes are involved in immunity, behavior, and other physiological pathways (Wittkopp and Beldade 2009). It is possible that some of the mutations characterized here also have additional, unscored physiological consequences in *Plodia*. Future work directly assessing immune function, behavior, or fitness-related traits in these mutants would help clarify the broader phenotypic effects.

Second, part of this study relies on the allele parsing capability of EditCo ICE v4, which infers CRISPR editing outcomes based on Sanger sequencing chromatograms through regression against candidate indel sequences (Conant et al. 2022). Unlike Nanopore or other next-generation sequencing approaches, this method does not directly sequence and count individual DNA molecules. Its estimates may therefore be less precise in highly complex samples, particularly in highly mosaic G_0_ individuals. Thus, ICE-based genotyping should be interpreted as evidence of editing and an approximate estimate of in/del composition rather than a definitive characterization of allelic variants present in a given sample. Accordingly, our study did not aim to establish which specific mutations or alleles were responsible for each observed phenotype. Future investigations at the genomic, transcriptomic, regulatory, and protein level would help clarify these genotype-phenotype relationships.

Third, we did not cover the pteridine pathway in *Plodia* pigmentation or its potential interactions with melanin and ommochrome pathways. Given that pteridines contribute to eye and cuticle coloration in several insects (Andrade and Carneiro 2021; Ferré 2024), some of the phenotypes observed in our mutants may also be influenced by pteridine pathway activity that was not directly assessed in this study. The potential of genes in this pathway as co-CRISPR markers also remain unexplored. *Plodia* does have orthologs of the *D. melanogaster* genes *Punch* (*Pi_Punch*, NCBI GeneID:128681158), *purple* (*Pi_purple*, NCBI GeneID:128676645), and *rosy* (*Pi_rosy*, NCBI GeneID:128676622), which are involved in the canonical pteridine pathway. These genes may offer complementary or more convenient screening options depending on the developmental stage or tissue of interest.

Finally, the co-CRISPR selection marker primarily serves as a visual proxy for editing at the GOI in the same G_0_ individual and does not guarantee that the GOI mutation is present in the germline or transmitted to the next generation. Because the marker gene and the GOI are typically located at unlinked loci, their mutant alleles generally assort independently during meiosis and are not necessarily inherited together by G_1_ offspring. Therefore, G_1_ and subsequent generations must be genotyped to confirm the presence and transmission of GOI mutations.

## MATERIALS AND METHODS

### Amino acid sequence alignment

The protein sequences in this study were obtained from Flybase *(Öztürk-Çolak et al. 2024)* and served as the query for an NCBI tBLASTn search to identify orthologs of *Plodia* interpunctella (Family: Pyralidae). Additional orthologs from related Lepidoptera species *Plutella xylostella* (Family: Plutellidae)*, Bombyx mori* (Family: Bombycidae)*, Spodoptera frugiperda* (Family: Noctuidae)*, Galleria mellonella* (Family: Pyralidae), and *Manduca sexta* (Family: Sphingidae), along with dipteran outgroup *Drosophila melanogaster* were identified using NCBI BLASTp. For each organism, the top seven sequences were selected based on query coverage, percent identity, and E-value, and imported into Geneious software as complete sequences (Geneious Prime 2025.1.2). Multiple sequence alignment was performed using Clustal Omega (v1.2.3).airwise protein sequence similarity was evaluated using the BLOSUM90 (BLOcks SUbstitution Matrix), a stringent scoring matrix for homologs, recently-diverged, or closely-related proteins, with threshold ≥ 0.

Geneious software processed % similarity by this formula:

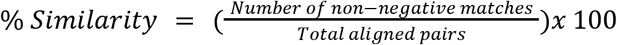

The number of non-negative matches are any two aligned amino acids that score ≥0 in BLOSUM90 chart. Total aligned pairs are every pair of compared amino acids across the alignment. The gaps are ignored.

### Plodia strain and husbandry

The *Plodia* strain used in this study is brown-eyed *bFog* strain (Heryanto et al. 2022a), a kind gift from Dr. Arnaud Martin (The George Washington University, Washington, DC). In our lab, this strain was reared at 28-29°C with controlled ambient humidity between 40-70% and 12:12 h light:dark photoperiod. The coarse wheat bran used in the *Plodia* diet was sterilized by autoclave. To reduce stickiness, we modified the diet recipe by reducing glycerol content to 24% w/w and adding 5% more water. Apart from these changes, the general husbandry and microinjection follows the The *Plodia* Manual made available through Open Science Framework (Heryanto et al. 2025).

### Embryonic microinjection

After the stock reached synchronized adulthood at ∼29 days post-fertilization, the container was exposed to CO_2_ gas to induce narcosis and oviposition. These adults were collected in the mason jar and kept at 28-29°C for the entire injection session (except during egg collection), as temperatures lower than RT reduced oviposition. Cas9 (QB3 MacroLab, UC Berkeley) and sgRNA (Synthego, CA) were mixed at 2:1 ratio (500:250 ng/µl final concentration) and incubated at RT for 10 minutes to form complexes prior to injection. The complex was then injected into freshly-laid embryos at RT with FemtoJet 4i (Calibre Scientific, CA) fitted with griphead set 4 size 0 (Calibre Scientific, Cat. No. 5196082001) holding a borosilicate capillary needle (WorldPrecision Instruments, Cat. No. 18100F-3) pulled at 57.8°C (Narishige PC-100). The initial setup for air pressure is 2.7 psi (*P_injection_*), 0.25 s (*t_injection_*), and 0.60 (*P_compensation_*), then adjusted to needle opening and droplet size. All embryos were injected between 10-50 min AEL and sealed with cyanoacrylate glue (GH1200 Super Glue Gel) then reared as described in the Plodia Manual (Heryanto et al. 2025).

For co-injection of sgRNAs, freshly laid eggs up to 50 minutes old were injected with a final mixture of 500 ng/μL CRISPR/Cas9, 125 ng/μL *white* or *scarlet* sgRNA and 125 ng/ul *yellow* (or GOI) sgRNA.

### sgRNA design

To design sgRNAs targeting eye and scale pigmentation genes, we used CRISPOR (Concordet and Haeussler 2018) to assess predicted KO efficiency and off-targets in the genome based on the *Plodia* genome ilPloInte3.2 (PRJNA782555). We prioritize targets within (1) the early exons, (2) highly-conserved exons determined by tBLASTn against *Pi, Bm, Px, Gm*, and *Sf,* or (3) if multiple isoforms are present, their shared DNA region. sgRNA candidates with predicted off-targets up to three mismatches in the exonic region are excluded.

### Screening and imaging of G_0_ knockouts at larval, pupal, and adult stages

Whenever possible, L4–L5 larvae exhibiting KO phenotypes were separated into a small petri dish (⌀50 mm) to monitor subsequent pupal and adult phenotypes. L4-L5 larvae, P8-P9 pupae, and freshly-emerged adults were screened using Leica S9i (Leica Microsystems, IL) with an integrated camera. KO phenotypes were compared to uninjected *bFog* control and verified with sequencing after they reached adulthood. To image the larvae live, they were placed in a petri dish on top of a PCR cooling rack until they were immobilized, but no more than 5 minutes. Freshly-emerged adults were sampled immediately after eclosion, frozen for >10 minutes, and assessed for scale intactness before dissecting the wings for imaging and gonads for sequencing. All images were adjusted in Photoshop to reflect the actual color observed through the microscope eyepiece.

To compare the normalized intensity between WT, *black*, and *ebony* mKO and to correct for ambient light variation during experiment, larval images (n=1 per genotype) of thoracic segments T1–T3 (**Figure 1C**) were imported into FIJI software (Schindelin et al. 2012). For each larva, the grayscale of multiple sclerite regions was measured by recording the area, mean gray value (*GV_sclerites_*), and standard deviation (*SD_sclerites_*). The same measurements were taken for non-pigmented regions (*GV_non-pigmented_, SD_non-pigmented_*). To convert GV to intensity (I), max grayvalue 255 is subtracted by mean GV.

The normalized intensity (NI) is calculated by this following formula:

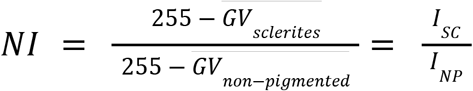

The standard deviation of this ratio was calculated using standard error propagation for division:

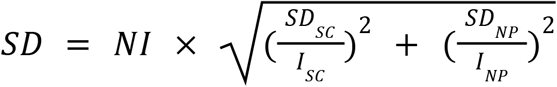

Similarly, the pupal sclerites (spiracles) measurements were compared to the rest of the puparium.

To show early melanization in *ebony* and *black* KO, early pupal stages (P2-P3) were selected based on the early eye pigment development (**Suppl. Fig. 3B, black arrow**). Since the acceleration of overall body pigment development coincides with the end of the eye pigmentation, WT pupae at stages just before (P6) and after (P9) eye pigment completion were used as comparison. The relative pigment intensity and SD of WT, *black*, and *ebony* marked in white dashes were directly plotted.

### Crossing

For in-crossing, G_0_ adults or late pupae with confirmed positive KO phenotype were mixed in a petri dish (⌀50mm). A piece of paper towel for climbing and a pinch of food for larvae were also added afterwards. After ∼5 days of matings, G_0_ adults were set aside for dissection and sequencing. All the crosses were placed in a small *Plodia* modified container with more food on the outside of the dish for the escapee. They were incubated at 29°C for about 1-2 weeks or until larvae were visible. At L5, cardboard cut-outs were added and the isolated pupae were scored for KO phenotype. G_1_ adults emerging from these pupae were then phenotyped and sequenced.

### Sanger Sequencing and ICE analysis

To extract DNA at G_0_, a small portion of adult female ovary with germarium or an adult male fused testes were submerged in DNARelease Buffer (Phire Tissue Direct PCR Kit), incubated for 5 minutes at RT then 2 minutes at 98°C. One microliter of the supernatant was used as a DNA template for PCR using Kapa HiFi HotStart ReadyMix (Roche Diagnostics, Cat. No. 07958935001) and primers **(Suppl. Table S6)** in a 25 µl reaction. Two microliters of PCR products were then used to run on 1% agarose electrophoresis gel in 1xTAE buffer.

To determine the DNA mutation at G_1_, the G_0_ adults with KO phenotype were firstly crossed randomly and the laid G_1_ eggs were reared into adulthood. An adult head of a G_1_ mutant individual (except for G_1_ *ebony* KO, a larval head) or a *Plodia bFog* WT control were then dissected for tissue-direct extraction for PCR reaction as mentioned above. Gel electrophoresis was done to confirm the presence of amplicons.

The PCR products were then purified with NucleoSpin Gel and PCR Clean-up Kit (Macherey Nagel, Cat. No. 740609.250). These samples were sent for sequencing and to parse the alleles, the Sanger chromatogram trace files (.ab1) of the mutants and WT were uploaded into a web-based software, Inference of CRISPR Edits (EditCo ICE v4) (Conant et al. 2022) as a batch. The alleles’ percentage contribution were categorized into in-frame, frameshift, WT, others, or large deletion. Large deletion was confirmed by comparing the amplicon size of the mutant against WT: (1) in gel electrophoresis result and (2) using Geneious software. These numbers were plotted using GraphPad Prism 10.

### Co-CRISPR performance analysis

The predictive performance of the co-CRISPR markers was evaluated using a 2x2 contingency table comparing marker phenotype with [yellow] phenotype. False positive rate was calculated as the proportion of marker-positive individuals that display WT phenotype (FP/(FP+TN)). False negative rate was calculated as the proportion of marker-negative individuals that display [yellow] phenotype (FN/(TP/FN)). Positive predictive value (PPV) was calculated as the proportion of marker-positive individuals that display [yellow] phenotype (TP/(TP+FP)). Negative predictive value (NPV) was calculated as the proportion of marker-negative individuals that display WT phenotype (TN/(TN+FN)).

For independent validation using meiosis-related genes, the PPV of [white] phenotype was calculated as the proportion of [white] individuals carrying an in/del at GOI locus, confirmed by Sanger sequencing and allele parsing.

## Data availability

All data supporting the conclusions of this study are presented in the article and its supplementary material.

## Acknowledgement

We thank Dr. Arnaud Martin (The George Washington University) for *Plodia interpunctella bFog, w^-/-^,* and *st^-/-^* strains, *scarlet* guide RNA, and for critical reading of the manuscript. We would also like to thank Dr. LingSze Lee and Dr. Selina Kindelay for critical reading of the manuscript, and Dr. Leif Benner for discussions at the early stages of the project. All schematics are made using Biorender.

## Author contributions

C.H. was responsible for conceptualization, experimentation and methodology, data analysis, writing, and editing. L.F.R. was responsible for conceptualization, editing, and supervision.

## References

Alqassar, J. D., Biot, M., Eccles, L. E., Stoppel, W. L., and Martin, A., 2025. Regionalization of gene expression and cell types in the silk glands of Plodia pantry moths. iScience 28: 11.

Andersen, S. O., 2010. Insect cuticular sclerotization: a review. Insect Biochem. Mol. Biol. 40: 166–178.

Andrade, P., and Carneiro, M., 2021. Pterin-based pigmentation in animals. Biol. Lett. 17: 20210221.

Bae, S., Kweon, J., Kim, H. S., and Kim, J.-S., 2014. Microhomology-based choice of Cas9 nuclease target sites. Nat. Methods 11: 705–706.

Banerjee, T. D., Finet, C., Seah, K. S., and Monteiro, A., 2024. Optix regulates nanomorphology of butterfly scales primarily via its effects on pigmentation. Front. Ecol. Evol. 12: 1392050.

Benner, L., Richmond, M., Xiang, Y., Lee, L., Hockens, C. B., Li, T., Yu, Z., Tsuchiya, D., Huang, S., Tromer, E. C., Hawley, R. S., and Rosin, L. F., 2026. Programmed meiotic errors facilitate dichotomous sperm production in the silkworm, Bombyx mori. Proc. Natl. Acad. Sci. 123: e2520991123.

Bi, H., Xu, J., He, L., Zhang, Y., Li, K., and Huang, Y., 2019. CRISPR/Cas9-mediated *ebony* knockout results in puparium melanism in *Spodoptera litura*. Insect Sci. 26: 1011–1019.

Binggeli, O., Neyen, C., Poidevin, M., and Lemaitre, B., 2014. Prophenoloxidase activation is required for survival to microbial infections in Drosophila. PLoS Pathog. 10: e1004067.

Borycz, J., Borycz, J. A., Loubani, M., and Meinertzhagen, I. A., 2002. Tan and ebony genes regulate a novel pathway for transmitter metabolism at fly photoreceptor terminals. J. Neurosci. 22: 10549–10557.

Camino, E. M., Butts, J. C., Ordway, A., Vellky, J. E., Rebeiz, M., and Williams, T. M., 2015. The evolutionary origination and diversification of a dimorphic gene regulatory network through parallel innovations in cis and trans. PLoS Genet. 11: e1005136.

Cheng, Y., Wang, P., Zeng, Y., An, W., Wang, T., and Xiao, Y., 2023. Characterization of five pigmentation genes as transgenic markers in Spodoptera frugiperda (Lepidoptera: Noctuidae). Int. J. Biol. Macromol. 242: 124981.

Conant, D., Hsiau, T., Rossi, N., Oki, J., Maures, T., Waite, K., Yang, J., Joshi, S., Kelso, R., Holden, K., Enzmann, B. L., and Stoner, R., 2022. Inference of CRISPR Edits from Sanger Trace Data. CRISPR J. 5: 123–130.

Concordet, J.-P., and Haeussler, M., 2018. CRISPOR: intuitive guide selection for CRISPR/Cas9 genome editing experiments and screens. Nucleic Acids Res. 46: W242–W245.

Dai, F., Qiao, L., Cao, C., Liu, X., Tong, X., He, S., Hu, H., Zhang, L., Wu, S., and Tan, D., 2015. Aspartate decarboxylase is required for a normal pupa pigmentation pattern in the silkworm, Bombyx mori. Sci. Rep. 5: 10885.

Dudzic, J. P., Kondo, S., Ueda, R., Bergman, C. M., and Lemaitre, B., 2015. Drosophila innate immunity: regional and functional specialization of prophenoloxidases. BMC Biol. 13: 81.

Eccles, L. E., Aikman, E. L., McTyer, J. B., Cruz, I. L. M., Richgels, A. L., and Stoppel, W. L., 2025. Exploring the functional properties of Plodia interpunctella silk fibers as a natural biopolymer for biomaterial applications. Mater. Today Commun. 42: 111416.

Eccles, L. E., Liwang, R. K., Orozco, A. A., Aikman, E. L., and Stoppel, W. L., 2026. Protein Sequence and Structural Diversity in *Plodia interpunctella* Silk Governs Microparticle Function and Macrophage Response. Biomacromolecules acs.biomac.6c00328.

Ewen-Campen, B., and Perrimon, N., 2018. ovoD Co-selection: A method for enriching CRISPR/Cas9-edited alleles in Drosophila. G3 Genes Genomes Genet. 8: 2749–2756.

Ferguson, L. C., and Jiggins, C. D., 2009. Shared and divergent expression domains on mimetic *Heliconius* wings. Evol. Dev. 11: 498–512.

Ferré, J., 2024. Biosynthesis of pteridines in insects: A review. Insects 15: 370.

Ferré, J., Silva, F. J., Real, M. D., and Ménsua, J. L., 1986. Pigment patterns in mutants affecting the biosynthesis of pteridines and xanthommatin in Drosophila melanogaster. Biochem. Genet. 24: 545–569.

Finkbeiner, S. D., Briscoe, A. D., and Reed, R. D., 2014. Warning signals are seductive: relative contributions of color and pattern to predator avoidance and mate attraction in Heliconius butterflies. Evolution 68: 3410–3420.

Fujiwara, H., and Nishikawa, H., 2016. Functional analysis of genes involved in color pattern formation in Lepidoptera. Curr. Opin. Insect Sci. 17: 16–23.

Futahashi, R., Koshikawa, S., Okude, G., and Osanai-Futahashi, M., 2022. Diversity of melanin synthesis genes in insects. Adv. Insect Physiol. 62: 339–376.

Ge, D. T., Tipping, C., Brodsky, M. H., and Zamore, P. D., 2016. Rapid screening for CRISPR-directed editing of the Drosophila genome using white coconversion. G3 Genes Genomes Genet. 6: 3197–3206.

Gilbert, L. E., Forrest, H. S., Schultz, T. D., and Harvey, D. J., 1988. Correlations of ultrastructure and pigmentation suggest how genes control development of wing scales of Heliconius butterflies.

Gonzalez-Baez, L., Mortati, E., Mitchell, L., and Losick, V. P., 2025. Melanization regulates wound healing by limiting polyploid cell growth in the Drosophila epithelium. Genetics 231: iyaf109.

Hanly, J. J., Loh, L. S., Mazo-Vargas, A., Rivera-Miranda, T. S., Livraghi, L., Tendolkar, A., Day, C. R., Liutikaite, N., Earls, E. A., and Corning, O. B., 2023. Frizzled2 receives WntA signaling during butterfly wing pattern formation. Development 150: dev201868.

Heryanto, C., Alqassar, J., Shodja, D., and Martin, A., 2025. A laboratory manual for the rearing, microinjection, and safe handling of the pantry moth Plodia interpunctella.

Heryanto, C., Hanly, J. J., Mazo-Vargas, A., Tendolkar, A., and Martin, A., 2022a. Mapping and CRISPR homology-directed repair of a recessive white eye mutation in Plodia moths. Iscience 25: 3.

Heryanto, C., Mazo-Vargas, A., and Martin, A., 2022b. Efficient hyperactive piggyBac transgenesis in Plodia pantry moths. Front. Genome Ed. 4: 1074888.

Hockens, C., Lorenzi, H., Wang, T. T., Lei, E. P., and Rosin, L. F., 2024. Chromosome segregation during spermatogenesis occurs through a unique center-kinetic mechanism in holocentric moth species. PLoS Genet. 20: e1011329.

Hong, J. W., Jeong, C. Y., Yu, J. H., Kim, S., Kang, S. K., Kim, S., Kim, N., Kim, K. Y., and Park, J. W., 2020. Bombyx mori kynurenine 3-monooxygenase gene editing and insect molecular breeding using the clustered regularly interspaced short palindromic repeat/CRISPR associated protein 9 system. Biotechnol. Prog. 36: e3054.

How, S. H. C., Banerjee, T. D., and Monteiro, A., 2023. Vermilion and cinnabar are involved in ommochrome pigment biosynthesis in eyes but not wings of Bicyclus anynana butterflies. Sci. Rep. 13: 9368.

Inoue, S., Watanabe, T., Hamaguchi, T., Ishimaru, Y., Miyawaki, K., Nikawa, T., Takahashi, A., Noji, S., and Mito, T., 2023. Combinatorial expression of ebony and tan generates body color variation from nymph through adult stages in the cricket, Gryllus bimaculatus. PLoS One 18: e0285934.

Ishizaki, Y., and Umebachi, Y., 1984. Synthesis, accumulation and excretion of kynurenine during the pupal and adult stages of Papilio xuthus. Insect Biochem. 14: 563–567.

Kim, H., Ishidate, T., Ghanta, K. S., Seth, M., Conte Jr, D., Shirayama, M., and Mello, C. C., 2014. A co-CRISPR strategy for efficient genome editing in Caenorhabditis elegans. Genetics 197: 1069–1080.

Klowden, M. J., 2013. Physiological systems in insects, Academic press.

Koch, P. B., 1991. Precursors of pattern specific ommatin in red wing scales of the polyphenic butterfly Araschnia levana L.: haemolymph tryptophan and 3-hydroxykynurenine. Insect Biochem. 21: 785–794.

Koch, P. B., Keys, D. N., Rocheleau, T., Aronstein, K., Blackburn, M., Carroll, S. B., and Constant, R. H. ffrench-, 1998. Regulation of dopa decarboxylase expression during colour pattern formation in wild-type and melanic tiger swallowtail butterflies. Development 125: 2303–2313.

Kolliopoulou, A., and Swevers, L., 2014. Recent progress in RNAi research in Lepidoptera: intracellular machinery, antiviral immune response and prospects for insect pest control. Curr. Opin. Insect Sci. 6: 28–34.

Kuwalekar, M., Deshmukh, R., Padvi, A., and Kunte, K., 2020. Molecular evolution and developmental expression of melanin pathway genes in Lepidoptera. Front. Ecol. Evol. 8: 226.

Lee, L., Mitra, A., Benner, L., Bruno, M., Raab, L. M., Runyan, R. K., Smith, E., Dale, R. K., and Rosin, L. F., 2026. Genome-wide crossover analysis in the holocentric pantry moth reveals strong crossover assurance and interference during spermatogenesis. Genome Res. 36: 1574–1589.

Li, J.-L., Li, S.-S., Luo, Z.-J., Lu, J., Cai, X.-M., Luo, Z.-X., Bian, L., Xiu, C.-L., Fu, N.-X., and Liu, N.-Y., 2024. CRISPR/Cas9-mediated ebony knockout causes melanin pigmentation and prevents moth Eclosion in Ectropis grisescens. Pestic. Biochem. Physiol. 200: 105810.

Li, X., Fan, D., Zhang, W., Liu, G., Zhang, L., Zhao, L., Fang, X., Chen, L., Dong, Y., and Chen, Y., 2015. Outbred genome sequencing and CRISPR/Cas9 gene editing in butterflies. Nat. Commun. 6: 8212.

Liao, C., Upadhyay, A., Liang, J., Han, Q., and Li, J., 2018. 3, 4-Dihydroxyphenylacetaldehyde synthase and cuticle formation in insects. Dev. Comp. Immunol. 83: 44–50.

Liu, Z., Zheng, W., Ye, W., Wang, C., Gao, Y., Cui, W., and Zhou, Z., 2019. Characterization of cysteine sulfinic acid decarboxylase from Tribolium castaneum and its application in the production of β-alanine. Appl. Microbiol. Biotechnol. 103: 9443–9453.

Livraghi, L., Hanly, J. J., Loh, L. S., Henry, A., Keck, C., Shirey, V. M., Tsai, C.-C., Yu, N., Van Belleghem, S. M., and Roberts, W. M., 2025. Genetic basis of an adaptive polymorphism controlling butterfly silver iridescence. Curr. Biol. 35: 2154–2163.

Loh, L. S., Hanly, J. J., Carter, A., Chatterjee, M., Tsimba, M., Shodja, D. N., Livraghi, L., Day, C. R., Reed, R. D., and McMillan, W. O., 2025. Single-nucleus transcriptomics of wing sexual dimorphism and scale cell specialization in sulphur butterflies. PLoS Biol. 23: e3003233.

Luo, L., Gui, S., Guo, Z., Feng, J., Smagghe, G., Liu, T., Liu, M., and Yi, T., 2025. Efficient CRISPR/Cas9-mediated *ebony* gene editing in the greater wax moth *Galleria mellonella*. Insect Sci. 32: 833–844.

Mackenzie, S. M., Brooker, M. R., Gill, T. R., Cox, G. B., Howells, A. J., and Ewart, G. D., 1999. Mutations in the white gene of Drosophila melanogaster affecting ABC transporters that determine eye colouration. Biochim. Biophys. Acta BBA-Biomembr. 1419: 173–185.

Massey, J. H., Chung, D., Siwanowicz, I., Stern, D. L., and Wittkopp, P. J., 2019. The yellow gene influences Drosophila male mating success through sex comb melanization. Elife 8: e49388.

Massey, J. H., and Wittkopp, P. J., 2016. The genetic basis of pigmentation differences within and between Drosophila species. Curr. Top. Dev. Biol. 119: 27–61.

Momose, T., De Cian, A., Shiba, K., Inaba, K., Giovannangeli, C., and Concordet, J.-P., 2018. High doses of CRISPR/Cas9 ribonucleoprotein efficiently induce gene knockout with low mosaicism in the hydrozoan Clytia hemisphaerica through microhomology-mediated deletion. Sci. Rep. 8: 11734.

Muth, F. W., 1969. Quantitative Untersuchungen zum Pigmentierungsgeschehen in den Augen der MehlmotteEphestia k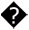hniella: III. Die Pigmentbildung im ?-Stamm nach. Injektion hoher Dosen von Kynurenin und der Versuch einer Deutung. Wilhelm Roux Arch. F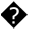r Entwicklungsmechanik Org. 162: 56–77.

Nakade, S., Tsubota, T., Sakane, Y., Kume, S., Sakamoto, N., Obara, M., Daimon, T., Sezutsu, H., Yamamoto, T., and Sakuma, T., 2014. Microhomology-mediated end-joining-dependent integration of donor DNA in cells and animals using TALENs and CRISPR/Cas9. Nat. Commun. 5: 5560.

Nie, H.-Y., Liang, L.-Q., Li, Q.-F., Li, Z.-H.-Q., Zhu, Y.-N., Guo, Y.-K., Zheng, Q.-L., Lin, Y., Yang, D.-L., and Li, Z.-G., 2021. CRISPR/Cas9 mediated knockout of Amyellow-y gene results in melanization defect of the cuticle in adult Apis mellifera. J. Insect Physiol. 132: 104264.

Okude, G., Futahashi, R., Kawahara-Miki, R., Yoshitake, K., Yajima, S., and Fukatsu, T., 2017. Electroporation-mediated RNA interference reveals a role of the multicopper oxidase 2 gene in dragonfly cuticular pigmentation. Appl. Entomol. Zool. 52: 379–387.

Öztürk-Çolak, A., Marygold, S. J., Antonazzo, G., Attrill, H., Goutte-Gattat, D., Jenkins, V. K., Matthews, B. B., Millburn, G., Dos Santos, G., and Tabone, C. J., 2024. FlyBase: updates to the Drosophila genes and genomes database. Genetics 227: iyad211.

Pearce, J. C., Campbell, J. S., Prior, J. L., Titball, R. W., and Wakefield, J. G., 2026. PiggyBac-mediated transgenesis and CRISPR–Cas9 knockout in the greater wax moth, Galleria mellonella. Lab Anim. 55: 95–102.

Quan, G.-X., Kim, I., Kômoto, N., Sezutsu, H., Ote, M., Shimada, T., Kanda, T., Mita, K., Kobayashi, M., and Tamura, T., 2002. Characterization of the kynurenine 3-monooxygenase gene corresponding to the white egg 1 mutant in the silkworm Bombyx mori. Mol. Genet. Genomics 267: 1–9.

Reed, R. D., McMillan, W. O., and Nagy, L. M., 2008. Gene expression underlying adaptive variation in Heliconius wing patterns: non-modular regulation of overlapping cinnabar and vermilion prepatterns. Proc. R. Soc. B Biol. Sci. 275: 37–46.

Reed, R. D., and Nagy, L. M., 2005. Evolutionary redeployment of a biosynthetic module: expression of eye pigment genes *vermilion*, *cinnabar*, and *white* in butterfly wing development. Evol. Dev. 7: 301–311.

Rogers, W. A., Grover, S., Stringer, S. J., Parks, J., Rebeiz, M., and Williams, T. M., 2014. A survey of the trans-regulatory landscape for Drosophila melanogaster abdominal pigmentation. Dev. Biol. 385: 417–432.

Romanowski, J. S., Myles, K. M., and Adelman, Z. N., 2026. Microhomology-mediated end joining is the predominant form of DNA repair in the mosquito Aedes aegypti with implications for gene editing, gene drive, and transgene removal. Nucleic Acids Res. 54: gkaf1532.

Ruxton, G. D., Allen, W. L., Sherratt, T. N., and Speed, M. P., 2019. Avoiding attack: the evolutionary ecology of crypsis, aposematism, and mimicry, Oxford university press.

Sakuma, T., Nakade, S., Sakane, Y., Suzuki, K.-I. T., and Yamamoto, T., 2016. MMEJ-assisted gene knock-in using TALENs and CRISPR-Cas9 with the PITCh systems. Nat. Protoc. 11: 118–133.

Schindelin, J., Arganda-Carreras, I., Frise, E., Kaynig, V., Longair, M., Pietzsch, T., Preibisch, S., Rueden, C., Saalfeld, S., and Schmid, B., 2012. Fiji: an open-source platform for biological-image analysis. Nat. Methods 9: 676–682.

Shirai, Y., Ohde, T., and Daimon, T., 2021. Functional conservation and diversification of yellow-y in lepidopteran insects. Insect Biochem. Mol. Biol. 128: 103515.

Shirk, B. D., Shirk, P. D., Furlong, R. B., Scully, E. D., Wu, K., and Siegfried, B. D., 2023. Gene editing of the ABC Transporter/White locus using CRISPR/Cas9-mediated mutagenesis in the Indian Meal Moth. J. Insect Physiol. 145: 104471.

Shodja, D. N., Livraghi, L., and Martin, A., 2025. Minos-mediated transgenesis in the pantry moth Plodia interpunctella. PeerJ 13: e20249.

Shodja, D. N., Yan, J., Livraghi, L., Yamada, A., Harvey-Samuel, T., Martin, A., and Mazo-Vargas, A., 2026. ABCG Transporter Knockouts Alter Integument and Eye Pigmentation With Gene-Specific Effects on Viability in Butterflies and Moths. Evol. Dev. 28: e70071.

Stevens, M., and Merilaita, S., 2008. Defining disruptive coloration and distinguishing its functions. Philos. Trans. R. Soc. B Biol. Sci. 364: 481.

Stratakis, E., 1982. Distribution and ontogeny of kynurenine 3-hydroxylase in Ephestia kühniella and its relation to metabolite levels. Insect Biochem. 12: 419–426.

Su, H., Yu, H., Xu, L., Zhang, M., Qi, Y., and Lu, Y., 2025. N-β-alanyl-dopamine synthetase gene (ebony) regulates pigmentation and reproduction in the melon fly, Zeugodacus cucurbitae. Pest Manag. Sci. 81: 3878–3888.

Sugumaran, M., and Barek, H., 2016. Critical analysis of the melanogenic pathway in insects and higher animals. Int. J. Mol. Sci. 17: 1753.

Sugumaran, M., and Lipke, H., 1982. Crosslink precursors for the dipteran puparium. Proc. Natl. Acad. Sci. 79: 2480–2484.

Terenius, O., Papanicolaou, A., Garbutt, J. S., Eleftherianos, I., Huvenne, H., Kanginakudru, S., Albrechtsen, M., An, C., Aymeric, J.-L., and Barthel, A., 2011. RNA interference in Lepidoptera: an overview of successful and unsuccessful studies and implications for experimental design. J. Insect Physiol. 57: 231–245.

Tomihara, K., Andolfatto, P., and Kiuchi, T., 2022. *Allele*-*specific* knockouts reveal a role for *apontic*-*like* in the evolutionary loss of larval melanin pigmentation in the domesticated silkworm, *Bombyx mori*. Insect Mol. Biol. 31: 701–710.

True, J. R., Yeh, S.-D., Hovemann, B. T., Kemme, T., Meinertzhagen, I. A., Edwards, T. N., Liou, S.-R., Han, Q., and Li, J., 2005. Drosophila tan encodes a novel hydrolase required in pigmentation and vision. PLoS Genet. 1: e63.

Uchiyama, H., Takasu, Y., Moriyama, M., Yoshitake, K., Uchiyama, H., Iizuka, T., Uchino, K., Okude, G., Banno, Y., and Kuwazaki, S., 2023. A novel monocarboxylate transporter involved in 3-hydroxykynurenine transport for ommochrome coloration. bioRxiv 2023–06.

Wang, Y., Huang, Y., Xu, X., Liu, Z., Li, J., Zhan, X., Yang, G., You, M., and You, S., 2021. CRISPR/Cas9-based functional analysis of yellow gene in the diamondback moth, *Plutella xylostella*. Insect Sci. 28: 1504–1509.

Wang, Y., Li, Z., Xu, J., Zeng, B., Ling, L., You, L., Chen, Y., Huang, Y., and Tan, A., 2013. The CRISPR/Cas system mediates efficient genome engineering in Bombyx mori. Cell Res. 23: 1414–1416.

Wei, W., Xin, H., Roy, B., Dai, J., Miao, Y., and Gao, G., 2014. Heritable genome editing with CRISPR/Cas9 in the silkworm, Bombyx mori. PloS One 9: e101210.

Wittkopp, P. J., and Beldade, P., 2009. Development and evolution of insect pigmentation: genetic mechanisms and the potential consequences of pleiotropy, pp. 65–71, Seminars in cell & developmental biology in Elsevier.

Wittkopp, P. J., Stewart, E. E., Arnold, L. L., Neidert, A. H., Haerum, B. K., Thompson, E. M., Akhras, S., Smith-Winberry, G., and Shefner, L., 2009. Intraspecific Polymorphism to Interspecific Divergence: Genetics of Pigmentation in *Drosophila*. Science 326: 540–544.

Wittkopp, P. J., True, J. R., and Carroll, S. B., 2002. Reciprocal functions of the Drosophila yellow and ebony proteins in the development and evolution of pigment patterns.

Xiang, Y., Tsuchiya, D., Guo, F., Gardner, J., McCroskey, S., Price, A., Tromer, E. C., Walters, J. R., Lake, C. M., and Hawley, R. S., 2023. A molecular cell biology toolkit for the study of meiosis in the silkworm Bombyx mori. G3 Genes Genomes Genet. 13: jkad058.

Xu, X., Harvey-Samuel, T., Yang, J., Alphey, L., and You, M., 2020. Ommochrome pathway genes kynurenine 3-hydroxylase and cardinal participate in eye pigmentation in Plutella xylostella. BMC Mol. Cell Biol. 21: 63.

Xu, X., Harvey-Samuel, T., Yang, J., You, M., and Alphey, L., 2021. CRISPR /Cas9-based functional characterization of the pigmentation gene *ebony* in *Plutella xylostella*. Insect Mol. Biol. 30: 615–623.

You, L., Bi, H., Wang, Y., Li, X., Chen, X., and Li, Z., 2019. CRISPR/Cas9-based mutation reveals *Argonaute 1* is essential for pigmentation in *Ostrinia furnacalis*. Insect Sci. 26: 1020–1028.

Zhang HaoKun, Z. H., Kiuchi, T., Hirayama, C., Katsuma, S., and Shimada, T., 2018. Bombyx ortholog of the Drosophila eye color gene brown controls riboflavin transport in Malpighian tubules.

Zhang, L., Martin, A., Perry, M. W., Burg, K. R. van der, Matsuoka, Y., Monteiro, A., and Reed, R. D., 2017a. Genetic basis of melanin pigmentation in butterfly wings. Genetics 205: 1537–1550.

Zhang, L., Mazo-Vargas, A., and Reed, R. D., 2017b. Single master regulatory gene coordinates the evolution and development of butterfly color and iridescence. Proc. Natl. Acad. Sci. 114: 10707–10712.

Zimlich, R. L., 2018. Moth Wing Nomenclature. BugGuide.

Zimowska, G., Silhacek, D. L., Shaaya, E., and Shirk, P. D., 1991. Immuno-Fluorescent analysis of follicular growth and development in whole ovaries of the Indianmeal moth. J. Morphol. 209: 215–228.

