## Supplementary Materials for "Pigmentation markers enable efficient co-CRISPR selection in the pantry moth *Plodia interpunctella*"

**Supplementary Table S1. Amino acid sequences used for pairwise comparison and multiple alignment.**

| Gene / Organism | ***black*** | ***ebony*** | ***yellow*** | ***tan*** | ***vermilion*** | ***kmo/cn*** | ***scarlet*** | ***white*** |
|---|---|---|---|---|---|---|---|---|
| *Plodia interpunctella* | XP_053606561.1 | XP_053619845.1 | XP_053600420.1 | XP_053623645.1 | XP_053616498.1 | XP_053625189.1 | XP_064293144.1 | NP_001037034.1 |
| *Plutella xylostella* | XP_011568918.3 | XP_026754130.2 | XP_011568334.3 | XP_011549615.3 | XP_037976399.2 | XP_048479642.1 | XP_048479461.1 | XP_053625431.1 |
| *Bombyx mori* | NP_001296491.1 | XP_030033133.2 | XP_062527627.1 | NP_001170882.1 | XP_004922659.1 | NP_001106135.1 | NP_001243922.1 | XP_026757709.2 |
| *Spodoptera frugiperda* | XP_035445371.2 | XP_050553942.1 | XP_035431942.2 | XP_035437652.1 | XP_050552036.1 | XP_035436851.2 | XP_050559103.1 | XP_011564267.2 |
| *Galleria mellonella* | XP_026762614.1 | XP_048486408.1 | XP_026765183.2 | XP_031767344.2 | XP_052749745.1 | XP_026757612.2 | XP_026757886.2 | XP_050559102.1 |
| *Manduca sexta* | XP_030035154.1 | XP_037876452.1 | XP_026765183.2 | XP_030022064.1 | XP_030023333.1 | XP_030033212.1 | XP_030034542.1 | XP_030034529.1 |
| *Drosophila melanogaster* | NP_001246025.1 | NP_524431.2 | NP_476792.1 | NP_001285042.1 | NP_511113.1 | NP_523651.4 | NP_524108.1 | NP_476787.1 |

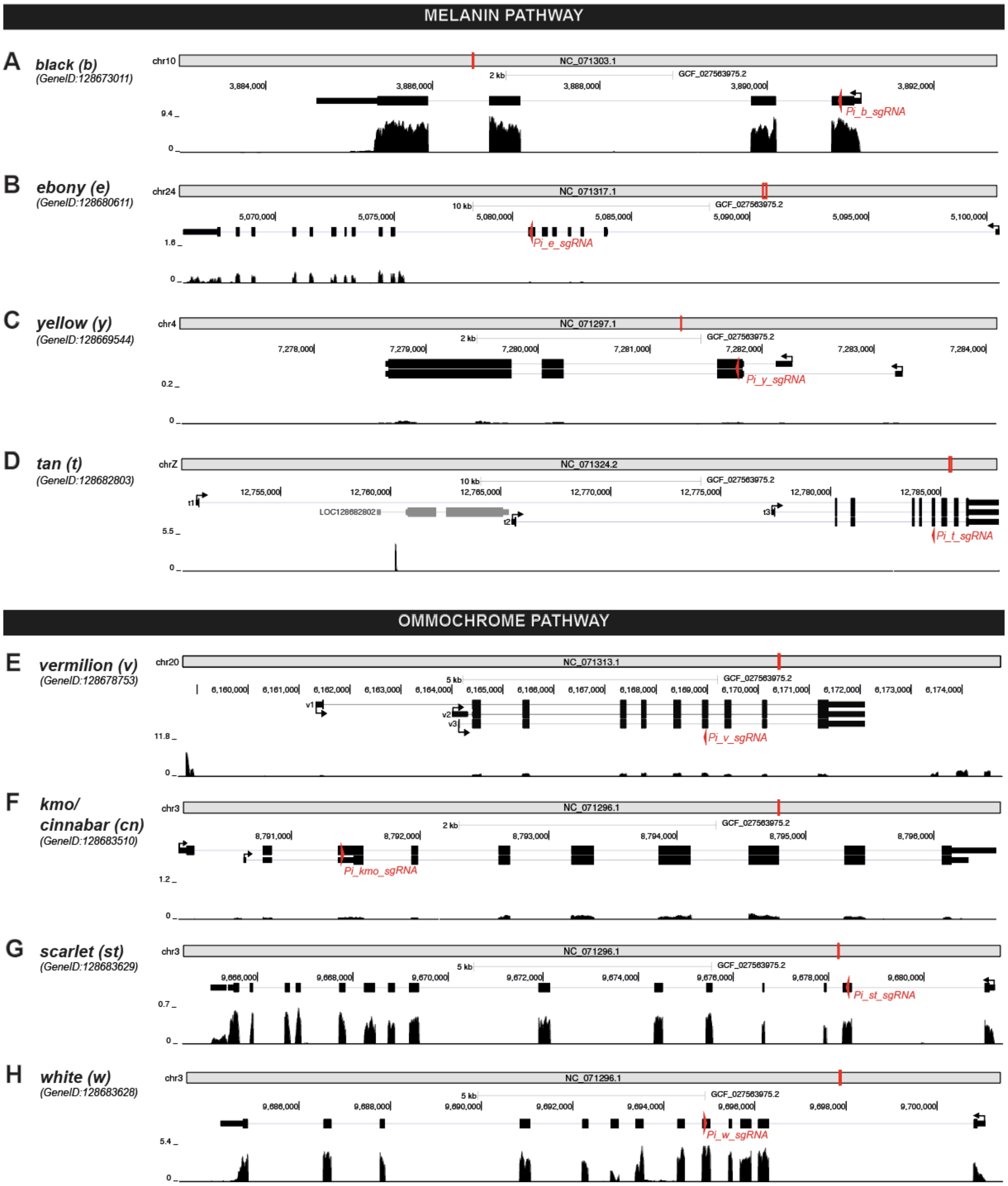

**Supplementary Figure S2. RNA-Seq of L5 *Plodia* testis on melanin and ommochrome pathway genes (Lee et al. 2026).**

RNA-Seq profile of (A) *black*, (B) *ebony*, (C) *yellow*, (D) *tan*, (E) *vermilion*, (F) *kmo/cinnabar*, (G) *scarlet*, and (H) *white* genes of the L5 *Plodia* testicular tissue. Red arrows denote the location and direction of the sgRNA target on the coding sequence.

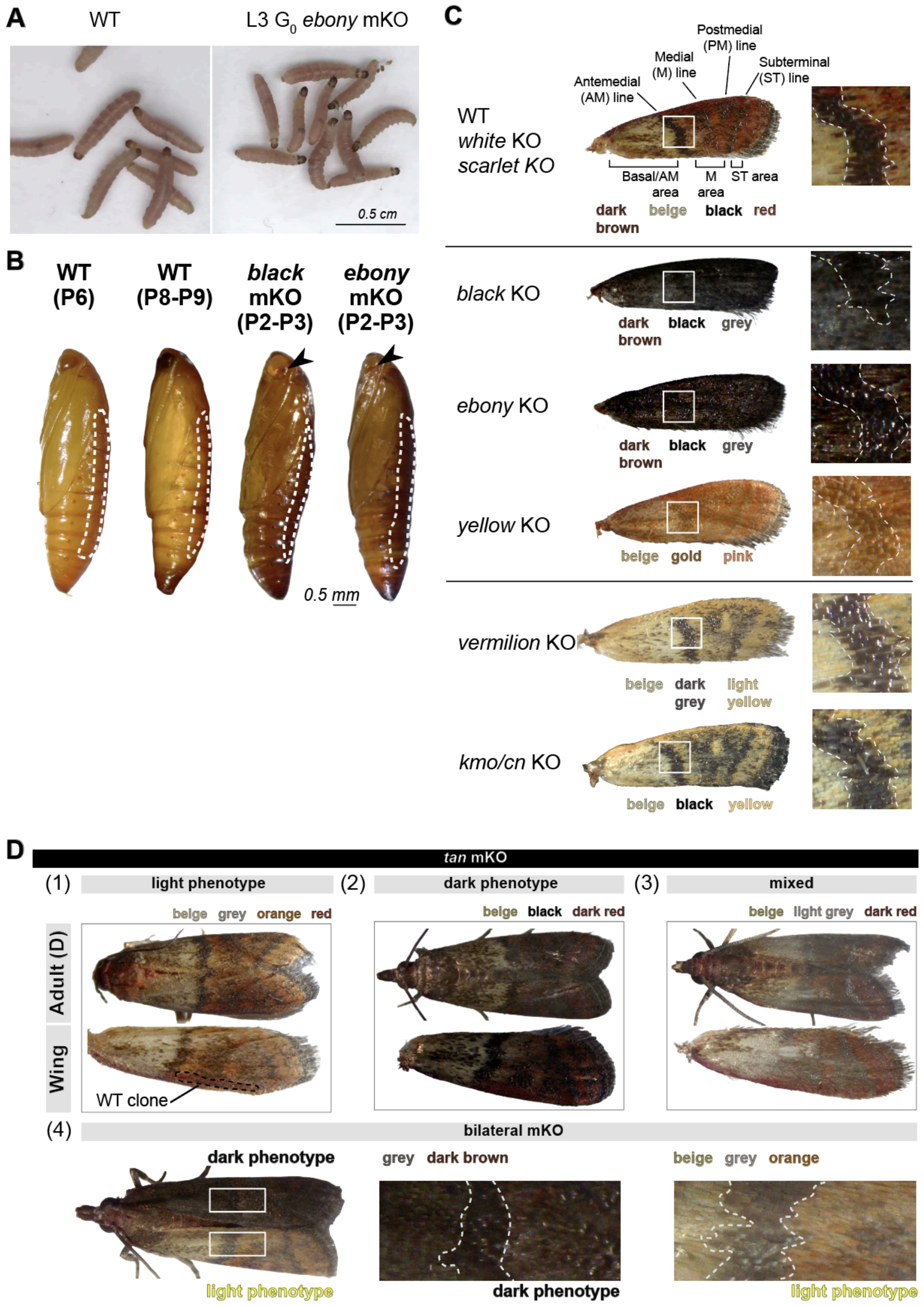

**Supplementary Figure S3.**

Raw image of (A) WT larvae (left) and *ebony* $G_0$ (right) at L3 larval stage taken with iPhone 15 Pro. The dark pigment of the head capsule and prothoracic shield is visible through the naked eye. This image is representative of the *black* $G_0$ larvae as well. (B) Puparium pigmentation. P6 and P8-P9 of WT pupae showed a stage before and after eye pigment deposition is completed, respectively. P2-P3 pupae of the *black* and *ebony* mKO showed body pigment has already begun well before eye pigment development is finished (black arrows). (C) Main color pigments on the forewing scales of *black, ebony, yellow, vermilion,* and *kmo/cn* mutants in comparison to WT color compartments. Insets show the area surrounding antemedial (AM) line, including AM area and medial (M) area. (D) Spectrum of *tan* mutant phenotypes and their main color pigments: (1) light phenotype, (2) dark phenotype, (3) mixed phenotype, and (4) bilateral phenotype.

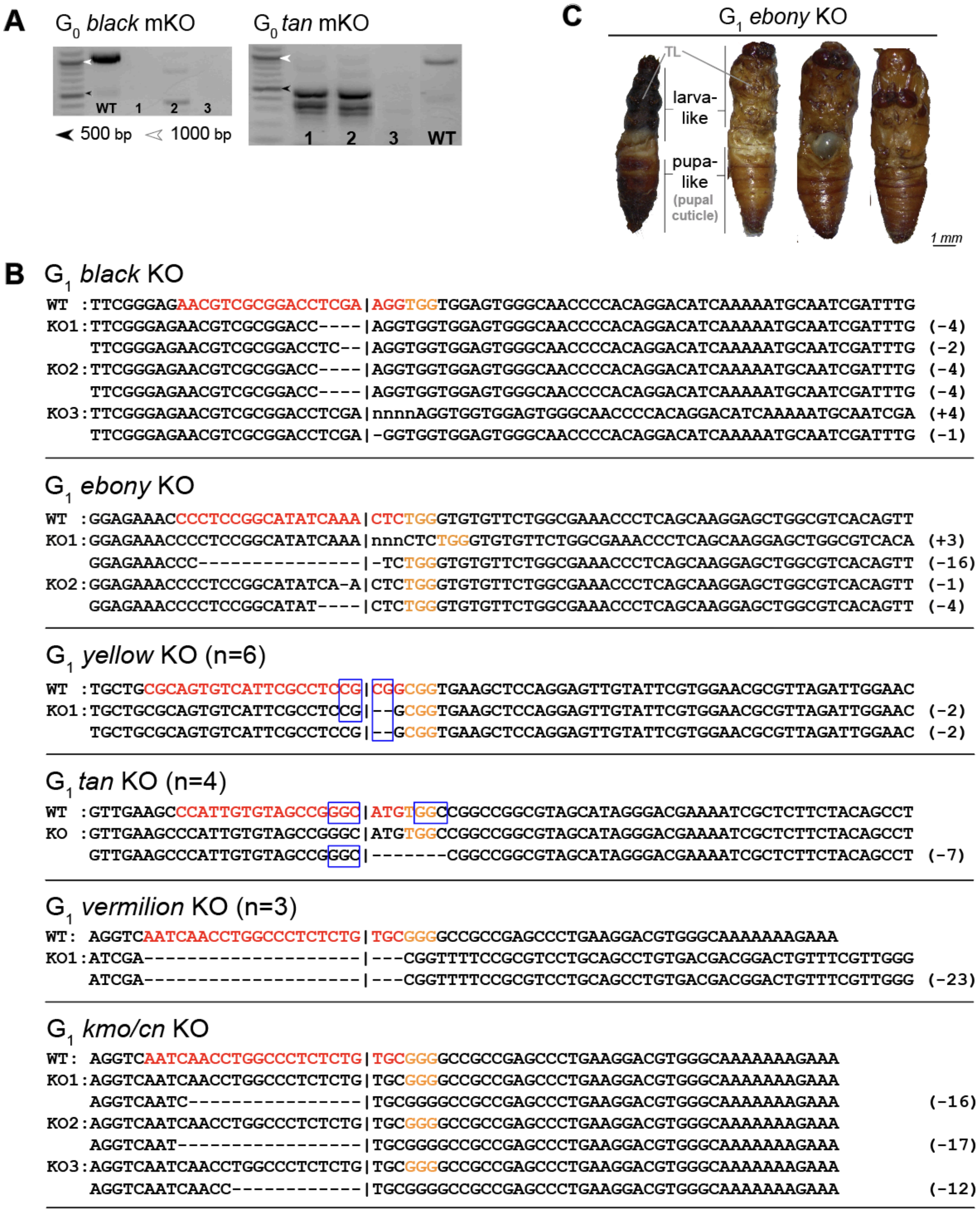

**Supplementary Figure S4.**

(A) Mutation analysis with gel electrophoresis of $G_0$ mKO gonadal tissues of *black* and *tan* mKO adults prior to Sanger sequencing. black arrow: 500 bp, white arrow: 1kb. (B) Some examples of sequenced $G_1$ mutant alleles from randomly-crossed $G_0$ crispants. Blue boxes: microhomology. Vertical bar: expected double-strand break (DSB) point. (C) Phenotype of $G_1$ *ebony* in-crosses. Failure in metamorphosis from L5 to pupal stage resulted in dead mutants with both half-larval and half pupal body.

**Supplementary Table S5. Summary of mutant phenotypes in this study.**

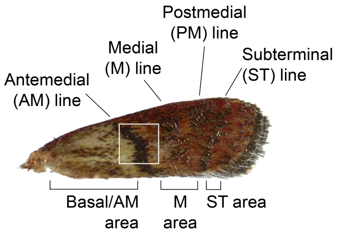

| Gene | Larval | | | Adult eyes | Adult wing pattern elements | | |
|---|---|---|---|---|---|---|---|
| | Sclerites | Cuticle | Testes | | Basal/AM area | Transverse lines | M, PM, and ST area |
| WT | light brown | pink to light red | red, brown | black | beige with brown spots | black | beige, red, brown |
| *melanin pathway* | | | | | | | |
| *black* | warm brown | red | red, warm brown | - | dark brown, gray | - | dark brown, gray |
| *ebony* | cool brown | red | cool brown | - | dark brown, gray | - | dark brown, gray |
| *yellow* | - | - | - | - | beige with pink spots | gold | beige, pink |
| *tan* | - | - | - | - | complete dark brown or beige, brown spots | dark gray | beige, orange or dark brown |
| *ommochrome pathway* | | | | | | | |
| *vermilion* | - | off-white | light yellow | orange, red | beige with light brown spots | light brown/gray | beige, light yellow |
| *kmo/cn* | - | off-white | yellow | yellow, red | beige with light brown spots | - | beige, yellow |
| *scarlet** | - | off-white | yellow | yellow | - | - | - |
| *white*** | - | off-white | translucent white | white | - | - | - |

*Transverse lines include antemedial (AM), medial (M), postmedial (PM), and subterminal (ST) lines (**Suppl. Fig. S3C**) (Zimlich 2018).

**Supplementary Table S6. CRISPR/Cas9 target sequences and genotyping primers used in this study**

| Gene | NCBI Gene ID | sgRNA | Target sequence, 5' to 3' | Sequencing Primer F | Sequencing Primer R |
|---|---|---|---|---|---|
| *melanin pathway* | | | | | |
| *black* | 128673011 | Pi_b_sgRNA | AACGTCGCGGACCTCGAAGG | GACTGGAGCCACTTCGTAGG | CAACAGTCCAAATGCCAGCC |
| *ebony* | 128680611 | Pi_e_sgRNA | CCCTCCGGCATATCAAACTC | TGGGAGGTCGTTGTTGATCC | AGCTACTCTACCGTCAGCCA |
| *yellow* | 128669544 | Pi_y_sgRNA | AGTGTCATTCGCCTCCGCGG | GTATACGCATTTTCAGCCACCG | CGTGGCATGACTGGACTTGG |
| *tan* | 128682803 | Pi_t_sgRNA | CCATTGTGTAGCCGGGCATG | TGTCTGCACACATCATCCCC | CTCGGGTGATGAACGTACGG |
| *ommochrome pathway* | | | | | |
| *vermilion* | 128678753 | *Pi_v_sgRNA* | TAGACCGGTAGATCTCGCGT | ACAGACGACATTGAGGCCG | TGATGAGGCTGTCGATGTCC |
| *kmo/cn* | 128683510 | Pi_kmo_sgRNA | CAACCTGGCCCTCTCTGTGC | TCTCGAAGCCCTTTACCTCG | GTTCTTCTAGGCCCACGTCC |
| *scarlet** | 128683629 | Pi_st_sgRNA | TGATTCCCTGCATCCCGAAG | TGTAATGACACGGTTATCGCA | GACCAGCGTCCCGGGTTTT |
| *white*** | 128683628 | *Pi_w_sgRNA* | TCGTCACAGAACATGAGTGG | CGAACATGGCGTATAACTCC | CCAGAACACAGTGATCGGC |

* (Shodja et al. 2026)
** (Heryanto et al. 2022a)

**Supplementary Table S7. *SPO11* sequence analysis**

| Label | Contribution | R Sq | Guide Sequences | Indels | Edited (Yes/No) |
|---|---|---|---|---|---|
| Control | 0 | 100 | GTTTCTGTCAGCAACGAATT | | |
| CVH105_spo11_27a | 0 | 100 | GTTTCTGTCAGCAACGAATT | | N |
| CH128_spo11_41 | 0 | 100 | GTTTCTGTCAGCAACGAATT | | N |
| CH124_spo11_33 | 3 | 99 | GTTTCTGTCAGCAACGAATT | {'-2': 2.0, '-1': 1.0} | Y |
| CH135_spo11_T | 10 | 10 | GTTTCTGTCAGCAACGAATT | {'-36': 2.0, '-6': 1.0, '-11': 2.0, '-31': 1.0, '-20': 1.0, '-32': 1.0, '-25': 1.0, '-4': 1.0} | Y |
| CH128_spo11_50 | 18 | 89 | GTTTCTGTCAGCAACGAATT | {'-22': 4.0, '2': 3.0, '-4': 3.0, '-16': 2.0, '-20': 1.0, '-1': 1.0, '17': 1.0, '7': 1.0, '-17': 1.0, '-19': 1.0} | Y |
| CVH136_spo11_FJ | 25 | 25 | GTTTCTGTCAGCAACGAATT | {'-1': 10.0, '1': 6.0, '-2': 3.0, '-31': 2.0, '-26': 2.0, '-29': 1.0, '-14': 1.0} | Y |
| CVH105_spo11_6 | 29 | 31 | GTTTCTGTCAGCAACGAATT | {'1': 3.0, '7': 3.0, '-15': 3.0, '20': 3.0, '-27': 2.0, '2': 2.0, '-11': 2.0, '-12': 2.0, '-31': 1.0, '-3': 1.0, '-16': 1.0, '-6': 1.0, '-25': 1.0, '4': 1.0, '5': 1.0, '11': 1.0, '15': 1.0} | Y |
| CVH105_spo11_2 | 34 | 94 | GTTTCTGTCAGCAACGAATT | {'7': 20.0, '-1': 14.0} | Y |
| CVH105_spo11_4 | 35 | 35 | GTTTCTGTCAGCAACGAATT | {'2': 8.0, '8': 4.0, '-10': 4.0, '-16': 4.0, '-14': 3.0, '-19': 2.0, '-11': 2.0, '6': 1.0, '-2': 1.0, '10': 1.0, '11': 1.0, '16': 1.0, '-13': 1.0, '17': 1.0, '-25': 1.0} | Y |
| CH135_spo11_K_A | 39 | 97 | GTTTCTGTCAGCAACGAATT | {'-4': 24.0, '-3': 11.0, '-1': 4.0} | Y |
| CVH105_spo11_5 | 45 | 92 | GTTTCTGTCAGCAACGAATT | {'-1': 24.0, '9': 10.0, '-4': 7.0, '-23': 1.0, '-3': 1.0, '1': 1.0, '-22': 1.0} | Y |
| CVH105_spo11_10 | 48 | 94 | GTTTCTGTCAGCAACGAATT | {'-1': 48.0} | Y |
| CH135_spo11_N | 51 | 92 | GTTTCTGTCAGCAACGAATT | {'-18': 44.0, '-1': 7.0} | Y |

| | | | | | |
|---|---|---|---|---|---|
| CH124_spo11_30 | 56 | 68 | GTTTCTGTCAGCAACGAATT | {'3': 32.0, '-2': 6.0, '-14': 3.0, '-5': 3.0, '12': 3.0, '-7': 2.0, '-25': 2.0, '-13': 2.0, '-12': 1.0, '-20': 1.0, '11': 1.0} | Y |
| CH124_spo11_32 | 61 | 61 | GTTTCTGTCAGCAACGAATT | {'-1': 35.0, '4': 6.0, '-19': 5.0, '17': 4.0, '-3': 4.0, '2': 2.0, '-26': 1.0, '-13': 1.0, '8': 1.0, '10': 1.0, '-9': 1.0} | Y |
| CH128_spo11_46 | 64 | 96 | GTTTCTGTCAGCAACGAATT | {'-1': 28.0, '-17': 33.0, '-27': 2.0, '6': 1.0} | Y |
| CH128_spo11_47 | 65 | 92 | GTTTCTGTCAGCAACGAATT | {'6': 31.0, '15': 23.0, '-4': 3.0, '1': 2.0, '-5': 2.0, '-2': 2.0, '-13': 1.0, '-16': 1.0} | Y |
| CH135_spo11_M | 66 | 66 | GTTTCTGTCAGCAACGAATT | {'3': 28.0, '-1': 7.0, '-30': 4.0, '-2': 5.0, '-29': 3.0, '-14': 3.0, '-8': 2.0, '-22': 2.0, '-17': 2.0, '-35': 1.0, '-19': 1.0, '-20': 1.0, '-9': 2.0, '-24': 1.0, '-26': 1.0, '-34': 1.0, '15': 1.0, '18': 1.0} | Y |
| CVH105_spo11_27 | 66 | 98 | GTTTCTGTCAGCAACGAATT | {'-1': 66.0} | Y |
| CH135_spo11_P | 69 | 95 | GTTTCTGTCAGCAACGAATT | {'4': 58.0, '-2': 5.0, '-5': 4.0, '-1': 2.0} | Y |
| CH124_spo11_36 | 71 | 71 | GTTTCTGTCAGCAACGAATT | {'10': 50.0, '-3': 5.0, '-24': 5.0, '-9': 3.0, '-25': 2.0, '-15': 2.0, '-10': 1.0, '-26': 1.0, '2': 1.0, '-21': 1.0} | Y |
| CH135_spo11_FG | 73 | 96 | GTTTCTGTCAGCAACGAATT | {'6': 24.0, '-4': 29.0, '-1': 20.0} | Y |
| CH128_spo11_37 | 76 | 88 | GTTTCTGTCAGCAACGAATT | {'10': 45.0, '6': 30.0, '-25': 1.0} | Y |
| CH128_spo11_38 | 76 | 96 | GTTTCTGTCAGCAACGAATT | {'6': 41.0, '-4': 34.0, '15': 1.0} | Y |

| | | | | | |
|---|---|---|---|---|---|
| CH135_spo11_L_A | 79 | 79 | GTTTCTGTCAGCAACGAATT | {'-4': 54.0, '-1': 8.0, '-2': 6.0, '11': 5.0, '-10': 2.0, '14': 2.0, '-9': 1.0, '8': 1.0} | Y |
| CH128_spo11_49 | 82 | 82 | GTTTCTGTCAGCAACGAATT | {'-18': 82.0} | Y |
| CH135_spo11_O | 84 | 98 | GTTTCTGTCAGCAACGAATT | {'-1': 45.0, '-4': 24.0, '9': 9.0, '7': 4.0, '6': 2.0} | Y |
| CH128_spo11_42 | 85 | 96 | GTTTCTGTCAGCAACGAATT | {'-1': 41.0, '4': 20.0, '2': 16.0, '1': 5.0, '-6': 2.0, '-4': 1.0} | Y |
| CH135_spo11_FF | 87 | 87 | GTTTCTGTCAGCAACGAATT | {'-1': 68.0, '1': 6.0, '19': 4.0, '10': 3.0, '-14': 1.0, '2': 1.0, '-26': 1.0, '18': 1.0, '-6': 1.0, '4': 1.0} | Y |
| CH124_spo11_C | 88 | 88 | GTTTCTGTCAGCAACGAATT | {'-32': 50.0, '5': 38.0} | Y |
| CH135_spo11_S | 88 | 88 | GTTTCTGTCAGCAACGAATT | {'9': 87.0, '-36': 1.0} | Y |
| CH124_spo11_B | 90 | 92 | GTTTCTGTCAGCAACGAATT | {'5': 44.0, '8': 39.0, '3': 4.0, '-3': 3.0} | Y |
| CH128_spo11_44 | 90 | 99 | GTTTCTGTCAGCAACGAATT | {'-6': 72.0, '-1': 11.0, '3': 4.0, '12': 3.0} | Y |
| CH128_spo11_53 | 90 | 90 | GTTTCTGTCAGCAACGAATT | {'3': 82.0, '-17': 4.0, '-32': 2.0, '-36': 1.0, '20': 1.0} | Y |
| CH124_spo11_A | 91 | 91 | GTTTCTGTCAGCAACGAATT | {'5': 48.0, '8': 43.0} | Y |
| CH124_spo11_31 | 91 | 91 | GTTTCTGTCAGCAACGAATT | {'-4': 49.0, '11': 42.0} | Y |
| CVH105_spo11_19 | 91 | 91 | GTTTCTGTCAGCAACGAATT | {'5': 48.0, '10': 43.0} | Y |
| CH128_spo11_54 | 91 | 94 | GTTTCTGTCAGCAACGAATT | {'-1': 77.0, '-3': 11.0, '3': 3.0} | Y |
| CVH105_spo11_24 | 92 | 92 | GTTTCTGTCAGCAACGAATT | {'-1': 48.0, '-24': 35.0, '-4': 5.0, '-3': 4.0} | Y |
| CVH117_spo11_12 | 92 | 92 | GTTTCTGTCAGCAACGAATT | {'7': 59.0, '-1': 33.0} | Y |
| CH135_spo11_V | 93 | 93 | GTTTCTGTCAGCAACGAATT | {'1': 46.0, '10': 39.0, '-1': 8.0} | Y |
| CVH105_spo11_13 | 93 | 93 | GTTTCTGTCAGCAACGAATT | {'7': 53.0, '1': 12.0, '-1': 21.0, '3': 7.0} | Y |
| CVH136_spo11_W | 94 | 94 | GTTTCTGTCAGCAACGAATT | {'6': 25.0, '-1': 34.0, '-4': 22.0, '4': 13.0} | Y |
| CH135_spo11_FE | 94 | 94 | GTTTCTGTCAGCAACGAATT | {'1': 51.0, '4': 38.0, '-3': 5.0} | Y |
| CVH105_spo11_8 | 94 | 94 | GTTTCTGTCAGCAACGAATT | {'5': 94.0} | Y |

| | | | | | |
|---|---|---|---|---|---|
| CVH105_spo11_14 | 94 | 94 | GTTTCTGTCAGCAACGAATT | {'-4': 33.0, '-16': 38.0, '9': 23.0} | Y |
| CVH105_spo11_20 | 94 | 94 | GTTTCTGTCAGCAACGAATT | {'3': 94.0} | Y |
| CVH105_spo11_25 | 94 | 94 | GTTTCTGTCAGCAACGAATT | {'5': 93.0, '-14': 1.0} | Y |
| CVH105_spo11_26 | 94 | 94 | GTTTCTGTCAGCAACGAATT | {'-17': 66.0, '11': 11.0, '16': 6.0, '-1': 7.0, '17': 4.0} | Y |
| CVH105_spo11_28 | 94 | 94 | GTTTCTGTCAGCAACGAATT | {'-1': 91.0, '4': 3.0} | Y |
| CH135_spo11_FI | 95 | 95 | GTTTCTGTCAGCAACGAATT | {'1': 27.0, '-2': 28.0, '2': 22.0, '-17': 13.0, '12': 5.0} | Y |
| CH128_spo11_45 | 95 | 95 | GTTTCTGTCAGCAACGAATT | {'1': 76.0, '-14': 8.0, '8': 6.0, '-25': 5.0} | Y |
| CH128_spo11_48 | 95 | 96 | GTTTCTGTCAGCAACGAATT | {'3': 63.0, '-1': 32.0} | Y |
| CVH136_spo11_AA | 96 | 96 | GTTTCTGTCAGCAACGAATT | {'4': 47.0, '-4': 49.0} | Y |
| CH135_spo11_FC | 96 | 96 | GTTTCTGTCAGCAACGAATT | {'-1': 56.0, '6': 40.0} | Y |
| CH124_spo11_34 | 96 | 96 | GTTTCTGTCAGCAACGAATT | {'-1': 80.0, '11': 6.0, '10': 4.0, '12': 3.0, '5': 2.0, '-4': 1.0} | Y |
| CVH105_spo11_15 | 96 | 96 | GTTTCTGTCAGCAACGAATT | {'2': 49.0, '-4': 47.0} | Y |
| CVH105_spo11_23 | 96 | 96 | GTTTCTGTCAGCAACGAATT | {'1': 56.0, '13': 21.0, '-1': 10.0, '-6': 9.0} | Y |
| CVH105_spo11_29 | 96 | 96 | GTTTCTGTCAGCAACGAATT | {'2': 90.0, '-1': 5.0, '-3': 1.0} | Y |
| CH124_spo11_D | 97 | 97 | GTTTCTGTCAGCAACGAATT | {'2': 71.0, '-1': 26.0} | Y |
| CH135_spo11_R | 97 | 97 | GTTTCTGTCAGCAACGAATT | {'-7': 76.0, '14': 21.0} | Y |
| CVH105_spo11_16 | 97 | 97 | GTTTCTGTCAGCAACGAATT | {'-1': 79.0, '1': 13.0, '-4': 5.0} | Y |
| CVH117_spo11_22 | 97 | 97 | GTTTCTGTCAGCAACGAATT | {'1': 55.0, '-4': 21.0, '-1': 21.0} | Y |
| CH128_spo11_39 | 97 | 97 | GTTTCTGTCAGCAACGAATT | {'-5': 60.0, '-2': 20.0, '3': 17.0} | Y |
| CH135_spo11_U | 98 | 98 | GTTTCTGTCAGCAACGAATT | {'-1': 56.0, '1': 22.0, '2': 20.0} | Y |
| CH135_spo11_FH | 98 | 98 | GTTTCTGTCAGCAACGAATT | {'-1': 73.0, '-4': 14.0, '-5': 11.0} | Y |
| CVH105_spo11_3 | 98 | 98 | GTTTCTGTCAGCAACGAATT | {'-1': 78.0, '4': 20.0} | Y |
| CVH105_spo11_21 | 98 | 98 | GTTTCTGTCAGCAACGAATT | {'-1': 98.0} | Y |
| CH128_spo11_40 | 98 | 98 | GTTTCTGTCAGCAACGAATT | {'2': 98.0} | Y |
| CH135_spo11_I_A | 99 | 99 | GTTTCTGTCAGCAACGAATT | {'-1': 99.0} | Y |

| | | | | | |
|---|---|---|---|---|---|
| CVH136_spo11_Z | 99 | 99 | GTTTCTGTCAGCAACGAATT | {'-1': 95.0, '-12': 4.0} | Y |
| CH135_spo11_FD | 99 | 99 | GTTTCTGTCAGCAACGAATT | {'-1': 83.0, '2': 16.0} | Y |
| CH135_spo11_J_A | 100 | 100 | GTTTCTGTCAGCAACGAATT | {'-1': 100.0} | Y |
| CVH136_spo11_X | 100 | 100 | GTTTCTGTCAGCAACGAATT | {'-1': 100.0} | Y |
| CH135_spo11_FB | 100 | 100 | GTTTCTGTCAGCAACGAATT | {'-1': 100.0} | Y |
| CVH136_spo11_FK | 100 | 100 | GTTTCTGTCAGCAACGAATT | {'-1': 100.0} | Y |
| CH124_spo11_35 | 100 | 100 | GTTTCTGTCAGCAACGAATT | {'-1': 100.0} | Y |
| CVH105_spo11_17 | 100 | 100 | GTTTCTGTCAGCAACGAATT | {'-1': 100.0} | Y |
| CH128_spo11_51 | 100 | 100 | GTTTCTGTCAGCAACGAATT | {'-5': 100.0} | Y |
| | | | | **Total Edited** | **77** |

**Supplementary Table S8.** ***SIX6OS1*** **sequence analysis**

| **Label** | **Contribution** | **R Sq** | **Guide Sequences** | **Indels** | **Edited (Yes/No)** |
|---|---|---|---|---|---|
| Control | 0 | 100 | CCTACCTTTAATGCAGTGGC | | |
| CVH127_six6os1_40 | 8 | 99 | CCTACCTTTAATGCAGTGGC | {'-2': 4.0, '-6': 4.0} | Y |
| six6os1_9 | 15 | 15 | CCTACCTTTAATGCAGTGGC | unparsable, traces manually checked | Y |
| six6os1_1 | 16 | 99 | CCTACCTTTAATGCAGTGGC | {'-6': 9.0, '-2': 5.0, '-3': 1.0, '-10': 1.0} | Y |
| CVH127_six6os1_30 | 16 | 99 | CCTACCTTTAATGCAGTGGC | {'-6': 9.0, '-3': 7.0} | Y |
| six6os1_7 | 17 | 98 | CCTACCTTTAATGCAGTGGC | {'-2': 7.0, '-6': 9.0, '-10': 1.0} | Y |
| CVH127_six6os1_27 | 18 | 99 | CCTACCTTTAATGCAGTGGC | {'-9': 4.0, '-2': 6.0, '-15': 3.0, '-10': 2.0, '-21': 3.0} | Y |
| CVH127_six6os1_25 | 30 | 99 | CCTACCTTTAATGCAGTGGC | {'-2': 17.0, '-3': 11.0, '-6': 1.0, '-9': 1.0} | Y |
| CVH127_six6os1_28 | 48 | 99 | CCTACCTTTAATGCAGTGGC | {'-2': 16.0, '-3': 18.0, '-6': 11.0, '-18': 1.0, '-9': 1.0, '-1': 1.0} | Y |
| CVH127_six6os1_42 | 72 | 93 | CCTACCTTTAATGCAGTGGC | {'-27': 33.0, '-2': 15.0, '-6': 15.0, '-9': 3.0, '20': 2.0, '-11': 2.0, '-8': 2.0} | Y |
| CVH127_six6os1_45 | 72 | 93 | CCTACCTTTAATGCAGTGGC | {'2': 49.0, '-16': 15.0, '10': 5.0, '-4': 3.0} | Y |

| | | | | | |
|---|---|---|---|---|---|
| CVH127_six6os1_23 | 75 | 84 | CCTACCTTTAATGCAGTGGC | {'10': 57.0, '-20': 6.0, '2': 3.0, '-3': 3.0, '-2': 2.0, '-11': 1.0, '-7': 1.0, '-12': 1.0, '-29': 1.0} | Y |
| CVH127_six6os1_44 | 84 | 92 | CCTACCTTTAATGCAGTGGC | {'1': 49.0, '8': 16.0, '-5': 6.0, '-6': 4.0, '-25': 4.0, '-15': 3.0, '-10': 1.0, '-21': 1.0} | Y |
| six6os1_21R | 85 | 97 | CCTACCTTTAATGCAGTGGC | {'-2': 26.0, '-6': 23.0, '3': 16.0, '1': 14.0, '-10': 5.0, '-15': 1.0} | Y |
| six6os1_5 | 86 | 86 | CCTACCTTTAATGCAGTGGC | {'14': 33.0, '-6': 19.0, '-2': 8.0, '-7': 7.0, '-5': 6.0, '-32': 3.0, '-14': 2.0, '-8': 3.0, '-11': 1.0, '-25': 1.0, '-27': 1.0, '11': 1.0, '-15': 1.0} | Y |
| six6os1_6 | 88 | 88 | CCTACCTTTAATGCAGTGGC | {'-3': 37.0, '-8': 25.0, '-7': 11.0, '-20': 5.0, '-15': 6.0, '-19': 2.0, '-33': 1.0, '-9': 1.0} | Y |
| CVH127_six6os1_46 | 89 | 89 | CCTACCTTTAATGCAGTGGC | {'4': 79.0, '-6': 10.0} | Y |
| CVH127_six6os1_29 | 90 | 99 | CCTACCTTTAATGCAGTGGC | {'-3': 57.0, '-2': 22.0, '-6': 5.0, '11': 3.0, '-9': 2.0, '-10': 1.0} | Y |
| six6os1_12 | 92 | 98 | CCTACCTTTAATGCAGTGGC | {'-6': 36.0, '2': 21.0, '-1': 20.0, '3': 5.0, '-2': 4.0, '-5': 6.0} | Y |
| six6os1_13 | 92 | 92 | CCTACCTTTAATGCAGTGGC | {'-16': 30.0, '4': 27.0, '-19': 31.0, '-10': 3.0, '-3': 1.0} | Y |
| six6os1_4 | 93 | 93 | CCTACCTTTAATGCAGTGGC | {'3': 51.0, '-6': 28.0, '-11': 4.0, '10': 4.0, '-3': 3.0, '4': 1.0, '-24': 1.0, '16': 1.0} | Y |
| six6os1_14 | 93 | 93 | CCTACCTTTAATGCAGTGGC | {'4': 48.0, '1': 28.0, '-19': 6.0, '-20': 3.0, '-14': 3.0, '-18': 3.0, '-25': 2.0} | Y |

| | | | | | |
|---|---|---|---|---|---|
| six6os1_22R | 93 | 95 | CCTACCTTTAATGCAGTGGC | {'-1': 19.0, '1': 17.0, '-2': 13.0, '3': 10.0, '-10': 24.0, '-6': 8.0, '-19': 1.0, '-20': 1.0} | Y |
| CVH127_six6os1_34 | 93 | 99 | CCTACCTTTAATGCAGTGGC | {'-6': 93.0} | Y |
| CVH127_six6os1_38 | 94 | 94 | CCTACCTTTAATGCAGTGGC | {'1': 81.0, '-2': 6.0, '-8': 4.0, '-6': 2.0, '-3': 1.0} | Y |
| six6os1_2 | 95 | 95 | CCTACCTTTAATGCAGTGGC | {'2': 47.0, '-4': 24.0, '-2': 22.0, '-11': 2.0} | Y |
| CVH127_six6os1_36 | 95 | 98 | CCTACCTTTAATGCAGTGGC | {'-2': 56.0, '2': 8.0, '1': 5.0, '7': 5.0, '-6': 7.0, '-1': 4.0, '-22': 4.0, '-3': 3.0, '-13': 2.0, '-9': 1.0} | Y |
| six6os1_16 | 96 | 96 | CCTACCTTTAATGCAGTGGC | {'-18': 45.0, '-2': 39.0, '15': 6.0, '-9': 6.0} | Y |
| CVH127_six6os1_31 | 97 | 98 | CCTACCTTTAATGCAGTGGC | {'-2': 59.0, '-3': 23.0, '-8': 11.0, '-9': 3.0, '4': 1.0} | Y |
| six6os1_8 | 98 | 98 | CCTACCTTTAATGCAGTGGC | {'-2': 98.0} | Y |
| six6os1_10 | 98 | 98 | CCTACCTTTAATGCAGTGGC | {'-2': 55.0, '-1': 34.0, '-6': 8.0, '-10': 1.0} | Y |
| six6os1_15 | 98 | 98 | CCTACCTTTAATGCAGTGGC | {'-6': 60.0, '-2': 19.0, '4': 11.0, '14': 8.0} | Y |
| six6os1_20 | 98 | 98 | CCTACCTTTAATGCAGTGGC | {'-2': 62.0, '-6': 17.0, '-13': 8.0, '1': 7.0, '-9': 4.0} | Y |
| CVH127_six6os1_41 | 98 | 98 | CCTACCTTTAATGCAGTGGC | {'-2': 79.0, '-27': 13.0, '-11': 2.0, '-8': 2.0, '-9': 1.0, '8': 1.0} | Y |
| six6os1_11 | 99 | 99 | CCTACCTTTAATGCAGTGGC | {'-3': 74.0, '11': 7.0, '-6': 8.0, '-2': 6.0, '-9': 2.0, '-5': 1.0, '3': 1.0} | Y |
| six6os1_17 | 99 | 99 | CCTACCTTTAATGCAGTGGC | {'-2': 63.0, '-3': 36.0} | Y |
| six6os1_18 | 99 | 99 | CCTACCTTTAATGCAGTGGC | {'-6': 99.0} | Y |
| CVH127_six6os1_39 | 99 | 99 | CCTACCTTTAATGCAGTGGC | {'-2': 95.0, '-6': 3.0, '-4': 1.0} | Y |
| CVH127_six6os1_43 | 99 | 99 | CCTACCTTTAATGCAGTGGC | {'-5': 92.0, '-2': 7.0} | Y |

| | | | | | |
|---|---|---|---|---|---|
| CVH127_six6os1_33 | | | CCTACCTTTAATGCAGTGGC | unparsable, traces manually checked | N |
| | | | | **Total Edited** | **38** |

**Supplementary Table S9. *SYCP3* sequence analysis**

| Label | Contribution | R Sq | Guide Sequences | Indels | Edited (Yes/No) |
|---|---|---|---|---|---|
| CVH115_15 | 0 | 100 | CGGAAAGCTGATACTCATGT | | |
| CVH115_20 | 0 | 100 | CGGAAAGCTGATACTCATGT | | N |
| CVH115_23 | 0 | 100 | CGGAAAGCTGATACTCATGT | | N |
| CVH115_12 | 5 | 98 | CGGAAAGCTGATACTCATGT | {'5': 5.0} | N |
| CVH115_24 | 5 | 98 | CGGAAAGCTGATACTCATGT | {'-1': 5.0} | N |
| CVH115_25 | 5 | 99 | CGGAAAGCTGATACTCATGT | {'5': 5.0} | N |
| CVH115_17 | 8 | 97 | CGGAAAGCTGATACTCATGT | {'1': 6.0, '3': 2.0} | N |
| CVH115_27 | 10 | 96 | CGGAAAGCTGATACTCATGT | {'-9': 7.0, '-2': 2.0, '-1': 1.0} | Y |
| CVH115_14 | 12 | 96 | CGGAAAGCTGATACTCATGT | {'-1': 6.0, '-16': 6.0} | Y |
| CVH115_26 | 12 | 95 | CGGAAAGCTGATACTCATGT | {'-28': 8.0, '-1': 2.0, '-2': 2.0} | Y |
| CVH115_11 | 17 | 96 | CGGAAAGCTGATACTCATGT | {'-2': 17.0} | Y |
| CVH107_y11 | 17 | 17 | CGGAAAGCTGATACTCATGT | {'-28': 9.0, '-23': 7.0, '18': 1.0} | Y |
| CVH115_3 | 21 | 89 | CGGAAAGCTGATACTCATGT | {'-21': 21.0} | Y |
| CVH115_22 | 25 | 80 | CGGAAAGCTGATACTCATGT | {'-1': 9.0, '-2': 16.0} | Y |
| CVH115_19 | 26 | 91 | CGGAAAGCTGATACTCATGT | {'-12': 10.0, '-15': 9.0, '-3': 4.0, '-2': 3.0} | Y |
| CVH115_5 | 27 | 82 | CGGAAAGCTGATACTCATGT | {'1': 27.0} | Y |
| CVH115_18 | 31 | 53 | CGGAAAGCTGATACTCATGT | unparsable, traces manually checked | Y |
| CVH115_2 | 34 | 78 | CGGAAAGCTGATACTCATGT | {'-4': 34.0} | Y |
| CVH115_10 | 35 | 79 | CGGAAAGCTGATACTCATGT | {'-3': 35.0} | Y |
| CVH115_4 | 40 | 82 | CGGAAAGCTGATACTCATGT | {'-6': 40.0} | Y |
| CVH115_1 | 55 | 55 | CGGAAAGCTGATACTCATGT | {'-1': 35.0, '8': 8.0, '-2': 12.0} | Y |
| CVH107_y4 | 78 | 84 | CGGAAAGCTGATACTCATGT | {'12': 41.0, '8': 21.0, '11': 9.0, '-1': 3.0, '3': 3.0, '-3': 1.0} | Y |

| | | | | | |
|---|---|---|---|---|---|
| CVH107_y10 | 82 | 89 | CGGAAAGCTGATACTCATGT | {'-6': 28.0, '-4': 28.0, '-9': 7.0, '4': 5.0, '-1': 6.0, '2': 4.0, '7': 2.0, '3': 1.0, '-3': 1.0} | Y |
| | | | | **Total Edited** | **16** |

**Supplementary Table S10. *HEI10* sequence analysis**

| Label | Contribution | R Sq | Guide Sequences | Indels | Edited (Yes/No) |
|---|---|---|---|---|---|
| Control | 0 | 100 | AAACATTTGGAGGCGCAGAT | | |
| CVH130_HEI10_n12 | 0 | 100 | AAACATTTGGAGGCGCAGAT | | N |
| CVH130_HEI10_n10 | 0 | 96 | AAACATTTGGAGGCGCAGAT | | N |
| CVH130_hei10_n20 | 11 | 11 | AAACATTTGGAGGCGCAGAT | {'-17': 4.0, '-37': 4.0, '-16': 2.0, '-39': 1.0} | Y |
| CVH130_HEI10_n11 | 11 | 11 | AAACATTTGGAGGCGCAGAT | {'-17': 2.0, '-24': 1.0, '-40': 1.0, '-11': 1.0, '-27': 1.0, '-26': 1.0, '-36': 1.0, '20': 1.0} | Y |
| CVH130_hei10_n36 | 12 | 70 | AAACATTTGGAGGCGCAGAT | {'1': 3.0, '-6': 2.0, '-16': 2.0, '-33': 2.0, '-11': 2.0, '-12': 1.0} | Y |
| CVH130_hei10_n26 | 13 | 14 | AAACATTTGGAGGCGCAGAT | unparsable, traces manually checked | Y |
| CVH130_HEI10_n14 | 14 | 97 | AAACATTTGGAGGCGCAGAT | {'1': 7.000000000000001, '-1': 6.0, '-5': 1.0} | Y |
| CVH130_hei10_n31 | 14 | 93 | AAACATTTGGAGGCGCAGAT | {'5': 14.000000000000002} | Y |
| CVH130_HEI10_n6 | 18 | 88 | AAACATTTGGAGGCGCAGAT | {'18': 13.0, '1': 5.0} | Y |
| CVH130_hei10_n22 | 18 | 93 | AAACATTTGGAGGCGCAGAT | {'5': 13.0, '7': 5.0} | Y |
| CVH130_HEI10_n19 | 22 | 87 | AAACATTTGGAGGCGCAGAT | {'14': 22.0} | Y |
| CVH130_HEI10_n15 | 23 | 23 | AAACATTTGGAGGCGCAGAT | {'3': 9.0, '15': 7.000000000000001, '2': 3.0, '-6': 2.0, '-30': 1.0, '-17': 1.0} | Y |

| | | | | | |
|---|---|---|---|---|---|
| CVH130_hei10_n22 b | 24 | 30 | AAACATTTGGAGGCGCAGAT | {'-1': 8.0, '5': 6.0, '6': 3.0, '-20': 2.0, '8': 2.0, '4': 1.0} | Y |
| CVH130_HEI10_n8 | 24 | 74 | AAACATTTGGAGGCGCAGAT | {'-11': 1.0, '-7': 6.0} | Y |
| CVH130_HEI10_n4 | 28 | 88 | AAACATTTGGAGGCGCAGAT | {'3': 22.0, '-11': 6.0} | Y |
| CVH130_HEI10_n17 | 29 | 90 | AAACATTTGGAGGCGCAGAT | {'1': 25.0, '-4': 4.0} | Y |
| CVH130_HEI10_n16 | 29 | 77 | AAACATTTGGAGGCGCAGAT | {'8': 25.0, '6': 3.0, '-4': 1.0} | Y |
| CVH130_hei10_n27 | 33 | 49 | AAACATTTGGAGGCGCAGAT | {'-1': 8.0, '15': 7.000000000000001, '-10': 2.0, '-32': 3.0, '-9': 2.0, '-29': 2.0, '14': 2.0, '-20': 1.0, '-5': 1.0, '-21': 1.0, '-17': 1.0} | Y |
| CVH130_HEI10_n5 | 35 | 86 | AAACATTTGGAGGCGCAGAT | {'-13': 2.0} | Y |
| CVH130_hei10_n25 | 37 | 64 | AAACATTTGGAGGCGCAGAT | {'-13': 9.0, '-6': 3.0, '-12': 2.0} | Y |
| CVH130_hei10_n29 | 37 | 74 | AAACATTTGGAGGCGCAGAT | {'5': 37.0} | Y |
| CVH130_hei10_n24 | 38 | 80 | AAACATTTGGAGGCGCAGAT | {'-1': 36.0, '5': 2.0} | Y |
| CVH130_hei10_n33 | 79 | 79 | AAACATTTGGAGGCGCAGAT | {'-40': 47.0, '-22': 1.0, '-21': 3.0, '-26': 1.0} | Y |
| CVH130_hei10_n21 | | | AAACATTTGGAGGCGCAGAT | unparsable, traces manually checked | Y |
| CVH130_HEI10_n3 | | | AAACATTTGGAGGCGCAGAT | unparsable, traces manually checked | Y |
| CVH130_HEI10_n9 | | | AAACATTTGGAGGCGCAGAT | unparsable, traces manually checked | Y |
| | | | | **Total Edited** | **24** |

**Supplementary Table S11.** ***DMC1*** **sequence analysis**

| **Label** | **Contribution** | **R Sq** | **Guide Sequences** | **Indels** | **Edited (Yes/No)** |
|---|---|---|---|---|---|
| CVH131_DMC1 | 1 | 95 | TCGGTCACTAACTTCTAGAG | {'-15': 1.0} | Y |
| CVH131_DMC1_d6 | 28 | 28 | TCGGTCACTAACTTCTAGAG | {'-17': 10.0, '-18': 7.0, '-25': 4.0, '-30': 3.0, '-20': 2.0, '-10': 2.0} | Y |
| CVH131_DMC1_d2 | 36 | 89 | TCGGTCACTAACTTCTAGAG | {'11': 15.0, '-4': 7.0, '-19': 12.0, '-14': 2.0} | Y |

| | | | | | |
|---|---|---|---|---|---|
| CVH131_DMC1_d16 | 52 | 86 | TCGGTCACTAACTTCTAGAG | {'-6': 44.0, '-12': 7.0, '-15': 1.0} | Y |
| CVH131_DMC1_d31 | 75 | 79 | TCGGTCACTAACTTCTAGAG | {'1': 23.0, '-2': 18.0, '-11': 10.0, '19': 6.0, '-27': 3.0, '-3': 3.0, '-12': 2.0, '-6': 3.0, '-13': 2.0, '6': 2.0, '-14': 1.0, '-9': 1.0, '-30': 1.0} | Y |
| CVH131_DMC1_d11 | 76 | 90 | TCGGTCACTAACTTCTAGAG | {'1': 34.0, '-2': 14.0, '-17': 20.0, '4': 2.0, '8': 2.0, '2': 2.0, '-18': 2.0} | Y |
| CVH131_DMC1_d23 | 80 | 80 | TCGGTCACTAACTTCTAGAG | {'5': 48.0, '-8': 17.0, '-4': 11.0, '-30': 2.0, '-25': 1.0, '-37': 1.0} | Y |
| CVH131_DMC1_d25 | 81 | 84 | TCGGTCACTAACTTCTAGAG | {'13': 25.0, '-3': 43.0, '-20': 6.0, '-22': 3.0, '-21': 2.0, '-30': 2.0} | Y |
| CVH131_DMC1_d15 | 84 | 84 | TCGGTCACTAACTTCTAGAG | {'-8': 18.0, '-1': 12.0, '-33': 12.0, '-2': 17.0, '9': 9.0, '-16': 3.0, '-19': 5.0, '-3': 3.0, '-17': 2.0, '10': 2.0, '5': 1.0} | Y |
| CVH131_DMC1_d12 | 84 | 84 | TCGGTCACTAACTTCTAGAG | {'3': 75.0, '-28': 3.0, '-10': 3.0, '-6': 3.0} | Y |
| CVH131_DMC1_d4 | 86 | 86 | TCGGTCACTAACTTCTAGAG | {'-20': 63.0, '-4': 11.0, '-6': 5.0, '-3': 3.0, '9': 3.0, '-29': 1.0} | Y |
| CVH131_DMC1_d22 | 86 | 87 | TCGGTCACTAACTTCTAGAG | {'1': 22.0, '9': 16.0, '-1': 12.0, '-20': 8.0, '4': 8.0, '-9': 6.0, '-30': 4.0, '-19': 6.0, '-6': 3.0, '-26': 1.0} | Y |
| CVH131_DMC1_d1 | 87 | 87 | TCGGTCACTAACTTCTAGAG | {'1': 37.0, '-7': 46.0, '-17': 2.0, '-25': 2.0} | Y |
| CVH131_DMC1_d14 | 88 | 89 | TCGGTCACTAACTTCTAGAG | {'4': 30.0, '5': 17.0, '1': 13.0, '-10': 14.0, '-11': 5.0, '-27': 2.0, '-29': 2.0, '3': 2.0, '-26': 1.0, '-7': 1.0, '-21': 1.0} | Y |
| CVH131_DMC1_d27 | 89 | 89 | TCGGTCACTAACTTCTAGAG | {'1': 83.0, '-18': 5.0, '-9': 1.0} | Y |

| | | | | | |
|---|---|---|---|---|---|
| CVH131_DMC1_d24 | 89 | 89 | TCGGTCACTAACTTCTAGAG | {'-3': 81.0, '-16': 6.0, '-25': 2.0} | Y |
| CVH131_DMC1_d21 | 89 | 89 | TCGGTCACTAACTTCTAGAG | {'-2': 40.0, '2': 25.0, '-33': 10.0, '-15': 5.0, '-21': 4.0, '-11': 3.0, '10': 1.0, '-16': 1.0} | Y |
| CVH131_DMC1_d18 | 90 | 90 | TCGGTCACTAACTTCTAGAG | {'-3': 58.0, '2': 30.0, '-12': 1.0, '-21': 1.0} | Y |
| CVH131_DMC1_d10 | 90 | 90 | TCGGTCACTAACTTCTAGAG | {'1': 73.0, '-4': 8.0, '-18': 6.0, '-8': 1.0, '-9': 2.0} | Y |
| CVH131_DMC1_d29 | 90 | 90 | TCGGTCACTAACTTCTAGAG | {'1': 34.0, '-2': 44.0, '-6': 7.0, '-18': 3.0, '-21': 2.0} | Y |
| CVH131_DMC1_d32 | 90 | 90 | TCGGTCACTAACTTCTAGAG | {'1': 82.0, '-18': 6.0, '-9': 2.0} | Y |
| CVH131_DMC1_d17 | 91 | 91 | TCGGTCACTAACTTCTAGAG | {'1': 38.0, '-2': 49.0, '-20': 3.0, '-21': 1.0} | Y |
| CVH131_DMC1_d8 | 91 | 91 | TCGGTCACTAACTTCTAGAG | {'1': 32.0, '3': 23.0, '-2': 33.0, '-22': 2.0, '-18': 1.0} | Y |
| CVH131_DMC1_d28 | 91 | 91 | TCGGTCACTAACTTCTAGAG | {'1': 54.0, '-6': 16.0, '-18': 7.0, '-14': 6.0, '-2': 3.0, '-3': 3.0, '3': 2.0} | Y |
| CVH131_DMC1_d20 | 92 | 92 | TCGGTCACTAACTTCTAGAG | {'-1': 43.0, '-6': 48.0, '-19': 1.0} | Y |
| CVH131_DMC1_d19 | 92 | 92 | TCGGTCACTAACTTCTAGAG | {'1': 50.0, '-2': 36.0, '-11': 3.0, '-15': 3.0} | Y |
| CVH131_DMC1_d13 | 92 | 94 | TCGGTCACTAACTTCTAGAG | {'-2': 74.0, '-21': 5.0, '-3': 8.0, '13': 2.0, '6': 1.0, '-4': 1.0, '-22': 1.0} | Y |
| CVH131_DMC1_d26 | 95 | 95 | TCGGTCACTAACTTCTAGAG | {'-2': 95.0} | Y |
| CVH131_DMC1_d9 | | | TCGGTCACTAACTTCTAGAG | unparsable, traces manually checked | N |
| | | | | **Total Edited** | **28** |

**Supplementary Table S12.** ***REC8*** **sequence analysis**

| Label | Contribution | R Sq | Guide Sequences | Indels | Edited (Yes/No) |
|---|---|---|---|---|---|

| | | | | | |
|---|---|---|---|---|---|
| Control | 0 | 100 | TATGCTGGTTAGCAGACTCG | | |
| CVH132_rec8_1 | 76 | 97 | TATGCTGGTTAGCAGACTCG | {'-1': 55.0, '-2': 21.0} | Y |
| CVH132_rec8_6 | 90 | 90 | TATGCTGGTTAGCAGACTCG | {'6': 62.0, '2': 11.0, '-1': 6.0, '7': 6.0, '-6': 3.0, '-3': 2.0} | Y |
| CVH132_rec8_8 | 76 | 95 | TATGCTGGTTAGCAGACTCG | {'-5': 53.0, '-3': 23.0} | Y |
| CVH132_rec8_11 | 42 | 89 | TATGCTGGTTAGCAGACTCG | {'6': 8.0, '-17': 12.0, '5': 5.0, '-2': 6.0, '16': 4.0, '-4': 2.0, '-12': 2.0, '1': 1.0, '-18': 1.0, '3': 1.0} | Y |
| CVH132_rec8_12 | 89 | 89 | TATGCTGGTTAGCAGACTCG | {'9': 38.0, '-1': 38.0, '20': 4.0, '-8': 4.0, '-9': 4.0, '11': 1.0} | Y |
| CVH132_rec8_10 | 90 | 90 | TATGCTGGTTAGCAGACTCG | {'3': 50.0, '-1': 26.0, '8': 14.0} | Y |
| CVH132_rec8_5 | 68 | 73 | TATGCTGGTTAGCAGACTCG | {'1': 14.0, '-3': 12.0, '-1': 12.0, '7': 6.0, '-16': 4.0, '-7': 3.0, '-5': 3.0, '20': 2.0, '11': 2.0, '-33': 2.0, '-11': 1.0, '-2': 1.0, '-20': 1.0, '-8': 1.0, '-19': 1.0, '-21': 1.0, '2': 1.0, '9': 1.0} | Y |
| CVH132_rec8_3 | 75 | 75 | TATGCTGGTTAGCAGACTCG | {'-3': 28.0, '-17': 43.0, '-7': 3.0, '-29': 1.0} | Y |
| | | | | **Total Edited** | **8** |
